# Biochemical analysis of anthocyanin and proanthocyanidin and their regulation in determining chickpea flower and seed coat colours

**DOI:** 10.1101/2022.08.22.504750

**Authors:** Lalita Pal, Vikas Dwivedi, Santosh Kumar Gupta, Samiksha Saxena, Ashutosh Pandey, Debasis Chattopadhyay

## Abstract

Flower and seed coat colour are important agronomic traits in chickpea (*Cicer arietinum* L.). Cultivated chickpeas are of two types namely, *desi* (dark seeded, purple flower) and *kabuli* (light colour seeded, white flower). There has been limited information about the molecular mechanism underlying the colour variation of flower and seed coats in *desi* and *kabuli* chickpea. We profiled the anthocyanin and proanthocyanidin (PA) contents in chickpea flowers and seed coats. Tissue-specific silencing of two genes encoding a basic helix-loop-helix (CabHLH) protein and a tonoplast-localized multidrug and toxic compound extrusion (CaMATE1) transporter in a *desi* genotype resulted in the reduction in expressions of anthocyanin and PA biosynthetic genes and anthocyanin and PA contents in the flower and seed coat and produced flowers and seeds with *kabuli* characteristics. Transcriptional regulation of a subset of anthocyanin and PA biosynthetic genes by a natural CabHLH variant and transport assay of a natural CaMATE1 variant explained the association of these alleles with the *kabuli* phenotype. We carried out a detailed molecular characterization of these genes, and provided evidences that *kabuli* chickpea flower and seed colour phenotype can be derived by manipulation of single genes in a *desi* chickpea background.

**Highlight:** In this study, we have defined the molecular link between flower and seed color in chickpea and identified CaMATE1 and CabHLH as the regulators of both the traits.

## Introduction

Chickpea (*Cicer arietinum* L.) is the second most important food grain legume crop in terms of production and consumption (FAOSTAT, 2020). Seed coat colour varies widely among cultivated chickpea genotypes and is considered an important agronomic trait as it influences consumer preference (Graham *et al*., 2001; Upadhyay and Ortz, 2001, Upadhyay *et al*., 2002; Hossain *et al*., 2010). Cultivated chickpea is categorized as ‘*desi*’ or ‘*kabuli*’ based on seed features such as colour and shape. Seeds of *desi* type chickpeas are generally dark brown in colour and angular in shape with a rough seed coat while the *kabuli* type produces light-brown coloured and rounded seeds with smooth seed coats (Upadhyay and Ortz, 2001, Upadhyay *et al*., 2002; Knights *et al*., 2010). Flower colour is a useful morphological marker in chickpea. The majority of the world’s chickpea germplasm accessions are pink-flowered or white-flowered. The dark-seeded *desi* types are typically associated with pink to purple coloured corolla and the *kabuli* types are characterized by white petals (Kumar *et al*., 2000; Hasan and Deb, 2013). Additional variations in these traits such as, blue flower and green seed coat occur at a very low frequency (Pundir *et al*., 1985).

Genetic regulations of flower and seed colours in chickpea have been studied for decades. A three-gene model with independently segregating genes, B, C and P had been proposed, where the homozygous recessive alleles of either B or C would confer white flower and the recessive homozygous allele of P is responsible for blue colour, whereas all three genes in dominant condition would produce pink colour (Ayyar and Balasubramanian, 1936; Kumar *et al*., 2000). However, the monogenic nature of inheritance of flower colour in some varieties has also been reported (Hasan and Deb, 2013). A genome-wide association study (GWAS) predicted fifteen major genomic loci exhibiting significant association with seed coat colour. The most significant one was mapped to a MATE (Multidrug and toxic compound extrusion) secondary transporter gene on chromosome 2 (Bajaj *et al*., 2015). The most important study in this aspect reported the identification of three deletions and two missense mutations in a basic helix-loop-helix (bHLH) transcription factor (TF) gene. Homozygous condition of any of these alleles was shown to be invariably associated with *kabuli* type chickpea (Penmetsa *et al*., 2016). Previously, induced mutation in the chickpea wild progenitor *C. reticulatum* directly produced kabuli phenotype suggesting that the flower and seed colours of chickpea might be monogenic and kabuli phenotype can be directly produced from *C. reticulatum* (Toker C, 2009).

Molecular mechanisms governing pigmentation in flowers and seeds have been characterized in many plant species. Many flavonoids, especially anthocyanins and proanthocyanidins (PAs), were detected as pigments that provide colour to the flower petals and seed coats (Lepiniec *et al*., 2006). Flavonoids are a large group of polyphenolic secondary metabolites biosynthesized from phenylalanine through multistep enzymatic processes. Their basic skeleton consists of two six-carbon aromatic rings connected by a three-carbon oxygenated heterocycle. Flavonoids are subclassified according to the oxidation state of the heterocycle and the compounds within the subclasses are characterized by the chemical modification of the side chains (Grotewold, 2006). PAs, also called condensed tannins, are the oligomers of flavon-3-ol derivatives. PAs such as oligomers of catechins and epicatechins are generally found in the seed coats and are responsible for seed coat pigmentation (Ariga *et al*., 1981; Gabetta *et al*., 2000; Gu *et al*., 2004; Dixon *et al*., 2005). The classical Mendelian study of genetic loci regulating floral pigmentation in garden pea defined two factors that conferred flower colour and regulates seed coat pigmentation (Hellens *et al*., 2010; Mendel, 1866; Moreau *et al*., 2012). One of those encodes a basic helix-loop-helix (bHLH) transcription factor (Hellens *et al*., 2010). The regulation of PA biosynthesis has been characterized by the analysis of *transparent testa (tt)* mutants that fail to accumulate PAs in the seed coat, making the seed coat transparent and revealing the yellow colour of the underlying cotyledons (Buerger 1971; Koornneef 1981, 1990). The role of bHLH family genes in anthocyanin and PA biosynthesis has been demonstrated in several plants. For example, in *Arabidopsis*, ENHANCER OF GLABRA3 (EGL3, a bHLH family protein) and PRODUCTION OF ANTHOCYANIN PIGMENT 1 (PAP1, a MYB family protein) interact to form MBW complex with TRANSPARENT TESTA GLABRA 1 (TTG1, a WD40 repeat family protein) to control the anthocyanin biosynthesis while, the ternary complex composed of TT8 (bHLH) and TT2 (MYB) and TTG1 transcriptionally regulate expression of the genes for PA biosynthesis (Baudry *et al*., 2004; Xu *et al*., 2013). In *Medicago truncatula*, MtTT8 interacts with MtTTG1 and different R2R3-MYB proteins MtLAP1 and MtPAR to regulate transcription of the late anthocyanin and PA biosynthesis genes, respectively (Peel *et al*., 2009; Verdier *et al*., 2012, Li *et al*., 2016). Although enzymes involved in flavonoid biosynthesis show various subcellular localizations, anthocyanins and PAs are commonly localized in vacuoles, where the monomeric units are polymerized and then produce brown oxidized products. Blocking flavonoid transport from the cytosol to central vacuole reduced anthocyanin and PA production, and genetic loss of function of a tonoplast-localized MATE1 transporter (an ortholog of Arabidopsis TT12) reduced PA biosynthesis in *M. truncatula* seeds (Abrahams *et al*., 2003; Goodman *et al*., 2004; Lepiniec *et al*., 2006; Zhao and Dixon, 2009).

Although the seed coat colour is an economically important trait of chickpea and the genome-wide association studies (GWAS) have predicted some genes and their alleles that are associated with this trait and the flower colour, the genes and their protein products were not biochemically characterized and there is no molecular evidence so far to show that these genes are responsible for seed coat colour. Earlier works indicated that AN1 (bHLH) of Petunia controls the colorations of both flower and seed coats (Chuck *et al*., 1993; Quatrocchio *et al*., 1993; Spelt *et al*., 2002). An 11-bp deletion in DFR gene resulted in a white flower morphology in *Iochroma calycinum* (Coburn *et al*., 2015). However, chemical basis for the seed phenotypes haven’t been reported in these cases. Further, there is no molecular evidence that links a single gene with the colours of both flower and seed coat of chickpea. Taking account of both the previous studies associating genes with these traits (Bajaj *et al*., 2015; Penmetsa *et al*., 2016), we developed knock-down lines of chickpea wherein the expression of orthologs of *MtTT8* and *MtMATE1* in chickpea (*CabHLH* and *CaMATE1*) were drastically reduced by tissue-specific RNA-interference (RNAi). We describe the molecular mechanisms underlying the transcriptional regulation of the anthocyanin and PA biosynthesis by the alleles of *CabHLH* and *CaMATE*1 governing chickpea flower and seed coat colours. Our results showed that downregulation of these two genes individually reduced anthocyanin and PA accumulation in flower and seed coat and altered their colours.

## Materials and Methods

### Plant materials and methods of chickpea transformation

Chickpea (*Cicer arietinum* L.) accessions IC296139 (var. Pusa-362) and ICC4958 for *desi* type and ICCV2, ICC95334 for *kabuli* type were used for all the studies. Genetic transformation of chickpea was performed with ICC4958 following an assimilated protocol from reported methods (Jayanand *et al*., 2003; Sharma *et al*., 2006; Chakaratborti *et al*., 2006; Bhatnagar-Mathur *et al*., 2009; Khandal *et al*., 2020). As reported before, a single cotyledon and half embryo was used as the explant (Khandal *et al*., 2020). Healthy matured seeds of ICC4958 were surface sterilized using sodium hypochlorite and soaked for 6-7 hrs in dark. Suspensions of *Agrobacterium tumefaciens* strain GV3101 bearing the relevant DNA constructs in buffer containing 1/2 MS (Murashige and Skoog medium, pH 5.5) and 100 μM acetosyringone were used for co-culturing with the explants for 30 min. The explants were incubated in co-culturing media for 48 hrs in dark at 22 °C. Explants having radicle were transferred to MS media containing cefotaxime (250 mg/L) and kanamycin (150 mg/L) for a few days for subculturing. Healthy shoots were used for micrografting on to seven-day-old soil-grown rootstocks of the same accession. The micrografted plants were grown in a growth chamber (Conviron, Winnipeg, Canada) at 22-24 °C with 60% humidity with a 10 hrs light period with an intensity of 250 µmol mt^-2^ sec^-1^ to maturity. The binary vectors *pBI101*.*2* and *pZP200* were used for raising promoter-β-glucuronidase (GUS) and RNAi transgenic lines, respectively. Positive transformants were detected by PCR using the specific primers for the intron used in the RNAi construct or the GUS-specific primers wherever applicable. All the primers sequences used in this study are mentioned in Supplementary Table S1.

### Description of RNAi and GUS constructs

A 332 bp-long intron of a *Brassica juncea* R2R3MYB gene (NCBI: JQ666168) was amplified from *B. juncea* genomic DNA with forward and reverse primer and cloned in *pGEMT*-T Easy plasmid (Invitrogen, Madison, WI). The sequence spanning the last 191 bp and 199 bp of the protein-coding regions of *CaMATE1*(*MtMATE1*) and *CabHLH* (*MtTT8*) genes, respectively, and the first 100 bp regions of their respective 3’-UTRs were used for the RNAi constructs. These gene fragments (*CaMATE1*:291 bp; *CabHLH*:299 bp) were amplified using the primers *MATE1*_SF and *MATE1*_SR; and *bHLH*_SF- and *bHLH*_SR primers and were cloned into pGEM-T Easy plasmid harbouring the *B. juncea* intron fragment using KpnI/SmaI and SacII restriction enzymes. The corresponding antisense DNA fragment was amplified with primers sets *MATE1*_AsF and *MATE1*_AsR; and *bHLH*_AsF and *bHLH*_AsR and cloned into pGEM-T Easy plasmid harbouring the *B. juncea* intron fragment and the corresponding sense DNA fragments using PstI and SacI. The full-length cassette was transferred into the *pZP200* plant binary vector using KpnI and SacI for *CaMATE1*, and SmaI and SacI for *CabHLH* RNAi cassette. A 1950 bp long *CaMATE1* promoter was amplified using primers *MATE1*p_F and *MATE1*p_R from ICC4958 chickpea DNA and cloned in *pZP200* vector harbouring the RNAi cassettes using XbaI and BamHI. To prepare promoter-reporter constructs 1950 bp and 1717 bp-long 5’-upstream sequences of *CaMATE1* and *CabHLH* genes were amplified from ICC4958 genomic DNA with the primers *MATE1*p_F/ *MATE*1p_R and *bHLH*p_F/ *bHLH*p_R and was cloned in *pBI101*.*2* plasmid using XbaI and BamHI.

### Histochemical staining

Tissues were vacuum infiltrated with GUS staining solution [50 mM sodium phosphate buffer (pH 7.0), 2 mM EDTA, 0.12% Triton, 0.4 mM ferrocyanide, 0.4 mM ferricyanide, 1.0 mM 5-bromo-4-chloro-3-indolyl-β-D-glucuronide cyclohexylammonium salt] for 20-30 min and incubated in the dark at 37 °C. The staining time varied between 2 days to 3 days depending on the tissue type. Tissues were cleared of chlorophyll by treatment with chloralhydrate at 75 °C for 2-3 hrs. The accumulation of PAs in the seed coat of chickpea was visualized by staining the whole seed with DMACA reagent (0.1% (w/v) DMACA in methanol-3 N HCL) for 25-30 min and then destained with ethanol:acetic acid (75:25, w/v) according to the method described previously (Zhao and Dixon, 2009). Stained seed coats were fixed with 4-5% agarose gel and dissected by microtome (Leica, VT1000 S, Wetzlar, Germany) and were visualized by Nikon 80i-epi-microscope. Ruthenium red solution (0.01%) was used to stain mucilage after imbibing seeds for 3-4 hrs with deionized water then washing 2-3 times with MilliQ water and dipping into the staining solution for 10 min (Pang *et al*., 2009).

### RNA Extraction and qRT-PCR

RNA was extracted from the plant tissues using Trizol (Merck, Branchburg, NJ) according to the manufacturer’s protocol. RNA from seed coats was extracted with GSure plant RNA isolation kit (GCC Biotech, Kolkata, India). Isolated RNA was treated with DNAse (Ambion, Elk Grove, CA). cDNA was synthesized from 1 μg of total RNA using oligodT and a High-fidelity cDNA synthesis kit (Thermo Scientific, Vilnius, Lithuania). The primers for quantitative real-time PCR (qRT-PCR) analysis are listed (Supplementary Table S1). Each qRT-PCR reaction was performed in 10 µL reaction volume containing appropriately diluted cDNA as template, 225 nM of each forward and reverse primer, and 2 X Power SYBR Green PCR master mix and Vii A 7 Real-Time PCR System (Applied Biosystem, Waltham, MA). Elongation factor 1-α (*CaEF-1*α) of chickpea (NM_001365163.1), the most suitable internal control (Garg *et al*., 2010) for normalizing gene expression in different tissues and developmental stages of chickpea, was used as the internal control for normalization. Uniform expression of *CaEF-1*α was checked with another internal control Actin (NCBI accession: AJ012685.1) (Supplementary Fig. S1A-B). Relative expression of genes was calculated according to delta-delta Ct method. Three independent biological replicates were used and three technical replicates were performed.

### Subcellular localization and BiFC

The open reading frame of both the *CaMATE1* and *CabHLH* was cloned in the gateway vector pENTER-D-Topo (Invitrogen, Waltham, MA) with primers MATE1pE_F, MATE1pE_R, and bHLHpE_F, bHLHpE_R, respectively (Supplementary Table S1) and then were transferred to the N-terminus of the *pEG101* binary vector to make GFP-fusion clones. *A. tumefaciens* strain EHA 105 harbouring *35S::CaMATE1-GFP* and *35S::CabHLH-GFP* constructs in infiltration media (10 mM MES-KOH, pH 5.6, 10 mM MgCl_2_, and 150 μM Acetosyringone) were infiltrated in *Nicotiana benthamiana* leaves for transient expression and viewed after 3 days under confocal microscope Leica TCS SP5 (Leica Microsystems) equipped with appropriate lasers (514 nm -527 nm and 532 nm-588 nm for YFP and RFP, respectively. The *Agrobacterium* cultures possessing organelle marker constructs for plasma membrane (CD3-1007 mCherry), vacuole (CD3-975 mCherry) (Nelson *et al*., 2007) and nucleus (CD3-1106 mCherry) (Citovsky *et al*., 2006) were mixed in 1:1 ratio with the *Agrobacterium* cultures of the experimental clones wherever required.

For bimolecular fluorescence complementation (BiFC), the full-length cDNA fragments (without the stop codon) were cloned into pENTER-D-TOPO by BP clonase reaction and then transferred to pSITE-nEYFPC1 (CD3-1648) or pSITEcEYFP-N1 (CD3-1651) plasmids (Martin *et al*., 2009). The *A. tumefaciens* strain EHA 105 harbouring the complementing BiFC constructs in 1:1 ratio were infiltrated as above in three-week-old *N. benthamiana* leaves and visualized after 60-65 hrs by the laser-scanning confocal microscope. Expression of proteins in the infiltrated samples was assessed by immunoblotting with anti-GFP antibody (Abcam, Cambridge, UK) following the standard protocol.

### Extraction and Quantification of PAs and Anthocyanins

Anthocyanin and PA were extracted and assayed following the published method (Pang *et al*., 2008). Briefly, tissues (0.25 to 1.0 g) were ground in liquid nitrogen and then extracted with 5 mL of extraction solution (70% acetone:0.5% acetic acid) by vortexing and sonication at 30 °C for 30 min. Following centrifugation at 2,500*g* for 10 min, residues were re-extracted twice as above. Pooled supernatants were then extracted with chloroform, and the aqueous supernatant was reextracted twice with chloroform and three times with hexane. Samples were freeze-dried and resuspended in extraction solution to a final concentration of 3 gm of original sample/ml. Soluble PA content was determined using DMACA reagent with catechin standard (Merck, Branchburg, NJ). Aliquots of samples or standards (2.5 μL) were mixed with 197.5 μL of DMACA reagent (0.2% [w/v] DMACA in methanol-3 N HCl [1:1]) in microplate wells. The absorbance was measured at wavelength 640 nm. For measurement of insoluble PAs, the residues from the above tissue extractions were dried in air for 2 days, and 1 mL butanol-HCl reagent was then added and the mixture sonicated at room temperature for 60 min, followed by centrifugation at 2,500*g* for 10 min (Pang *et al*., 2007). Absorbance at 550 nm of the supernatant was taken before and after boiling. The difference in absorbance values were converted into PA equivalents using a standard curve of procyanidin B1 (Merck, Branchburg, NJ).

For analysis of total anthocyanin content (TAC), 2 mL of acidified methanol (1% v/v) was added to 0.25 gm of ground tissue and the extract was incubated overnight at 4 °C in dark. The extract was centrifuged at 3000*g*. The absorbance of the supernatant was read at 530 and 657 nm. The relative unit was calculated using the following formula: (A530 - 0.25 * A657)/weight of tissue (Mishra *et al*., 2016).

### Ultra High Performance Liquid Chromatography-Mass spectrometry (LC-MS) analysis

A previously published method was followed (Naik *et al*., 2021). Dried lyophilized tissues (250 mg) were extracted with 80% methanol. The clear supernatant was sterilized through a 0.22 µm and used for anthocyanin estimation. Three volumes of 2 M Methanol:HCL was added and incubated at 90-95 °C for 50 min. Finally, the samples were dried off in a rotavapor and dissolved in 1mL of 80% methanol. Three biological replicates were analysed qualitatively and quantitatively by 1290 Infinity II series UHPLC system (Agilent Technologies, Santa Clara, CA) equipped with Zorbax Eclipse Plus C_18_ column maintained at 30 °C. The mobile phase consisted of an aqueous solution of 0.1% formic acid (solution A) and 0.1% formic acid in acetonitrile (solution B). The gradient for solution B was programmed as described previously. Other chromatographic parameters included a constant flow of 270 µL/min (injection volume, 3 µL), and a run time of 47 min including equilibration. Before analysis, all samples were filtered through a 0.22 µm PVDF syringe filter (Merck). Analysis of the proanthocyanidins was performed using a UHPLC system (Exion LC Sciex, Place) coupled to a triple quadrupole system (QTRAP6500+) (ABSciex, Place) using electrospray ionization. The voltage was set at 5,500V for positive ionization. The mass spectrometer was used in multiple reaction monitoring modes for qualitative and quantitative analysis by using Analyst software (version 1.5.2). Analytical standards (Merck, Branchburg, NJ) were used for each compound.

### Transactivation assay

Transactivation assays were performed with the effecter constructs (*CabHLH, CabHLHb2, CaPAR, CaLAP1*, and *CaWD40-1*) cloned in *pBTDest* vector for At7 *Arabidopsis* protoplast co-transfection assay. The reporter constructs include promoters of *C. aeritinum LDOX, BAN*, and *DFR* genes amplified from chickpea genomic DNA using specific primers and cloned to the *p-DISCO* vector, resulting in *Promoter*::*GUS*reporter constructs. At7 cell line protoplast (Stracke *et al*., 2016) was used for this assay. A purified 25 µg of premixed plasmid DNA having 10 µg of reporter construct, 10 µg of effector construct, and 5 µg of LUC plasmid (pBT-pUb14-2-Luc), and a filler plasmid (pBT10-Δ-LUC) to adjust the total plasmid amount (Sprenger-Haussels and Weisshaar, 2000) for transfection normalization were used in co-transfection into At7 cell line protoplast suspension. The transfected protoplasts were incubated for 20 h-22 hrs at 26 ºC in the dark followed by harvesting the protoplast in 96 well plates for luciferase, GUS (ex: 365 nm/em: 460 nm), and Bradford assays (595 nm). Normalized specific GUS activity, as a measure for TF activation capacity, was presented in pmol 4-methylumbelliferone (4-MU) mg^-1^ of protein min^-1^).

### Preparation of yeast membrane vesicle and transport activity assay

*Saccharomyces cerevisiae* AD12345678 strain (Morita *et al*., 2009) was transformed with *CaMATE1* cDNA and its mutated form cloned in *pDR196* yeast expression vector. The positive transformants were grown in YNB (-Uracil)+ 2% dextrose medium overnight at 30 °C. Yeast microsomal fraction was prepared following the previously reported procedure (Nakanishi *et al*., 2001). Briefly, the cells palette was digested with Zymolyase media-20T (MP Biomedical, OH) for 2 hrs at 30 °C with gentle agitation. The digested suspension containing spheroplasts were homogenised with acid-washed glass beads (0.5 mm). The cell debris were precipitated by centrifuge at 2000*g* for 10 min and the supernatant was used to precipitate the microsomal fraction at 120,000*g* for 30 min. The pellet was resuspended in the buffer (Tris-MES, pH 7.6, 0.3 M sorbitol, 1 mM DTT, 0.1 M KCl, and protease inhibitor cocktail). A transport activity assay was performed according to the method described in Bartholomew *et al*., 2002. 1 mL reaction buffer (25 mM Tris-MES pH-8, 0.4 M sorbitol, 50 mM KCl, 5 mM Mg-ATP, 0.1% (w/v) BSA) and 100 µM of Cyanidin 3’O-glucoside was mixed with 50 µg of microsomal fraction and incubated at 25 °C. A total of 100 µL of reaction was taken out at different time intervals and the reaction was stopped with ice-cold washing buffer (25 mM Tris-MES, pH-8.0, 0.4 M sorbitol). The reaction mixture was passed through 0.22 µM polyvinylidene difluoride (PVDF) membrane filters and the filters containing vesicles were eluted with 1 ml of 50% (v/v) methanol. The elute was analysed by LC-MS.

## Results

### Assessment of anthocyanin- and proanthocyanidin precursor contents in the flower and seed coat of chickpea

A *desi* type chickpea typically produces pink-to-purple flower petals and dark brown coloured seeds and the *kabuli* types are recognized by white petals and light-coloured seeds (Fig. 1A, 2A). Sections of these two types of seed coats were stained with 4-Dimethylaminocinnamaldehyde (DMACA). The inner integument and the endothelial layers of the *desi* seed showed intense deposition of DMACA-reactive compound (Proanthocyanidin) in contrast to the *kabuli* seed showing no detectable PA deposition (Fig. 1B). Contents of specific PA precursors, catechin, epigallocatechin, and epicatechin gallate were measured in the mature seed coats of two *desi* (IC296139, ICC4958) and two *kabuli* (ICCV2, ICC95334) (Fig. 1A) type chickpea accessions using liquid chromatography-coupled-mass spectrometry (LC-MS). Expectedly, both the *desi* types showed very high contents of these PA precursors, whereas those compounds were either below the detection limit or present in a negligible amount in seed coats of both the *kabuli* type chickpeas (Fig. 1C). Similarly, the anthocyanin precursors such as delphinidin, pelargonidin, cyanidin, malvidin, and peonidin contents were present in much higher levels in *desi* seeds in contrast to being undetectable levels in *kabuli* seed coats (Fig. 1D). In flower petals, the six anthocyanin precursors (cyanidin, delphinidin, petunidin, pelargonidin, peonidin and malvidin) were measured, but only delphinidin was detectable in the *desi* chickpea petals and the others were found to be present below detection limit. Beside this, no anthocyanin was present in detectable range in the *kabuli* flower petals (Fig. 2B). Levels of PA precursors were negligible in chickpea flower petals. Expression of the genes encoding leucoanthocyanin dioxygenase (*CaLDOX*) (LOC101511289) and BANYULS (*CaBAN*) (LOC101494604), two important enzymes in the flavonoid pathway are considered as the indicators of anthocyanin and proanthocyanidin production, respectively (Pelletier *et al*., 1997, Xie *et al*., 2003). The expression levels of these two genes were quantified in the petals and seed coats of these chickpea accessions. *CaLDOX* gene was predominantly expressed in the coloured petals, whereas the expression of *CaBAN* gene was almost absent even in the *desi* type petals. *CaLDOX* gene expression in the *kabuli* type chickpea petals was apparently absent in petals. In contrast, dark-coloured seed coats primarily exhibited *CaBAN* gene expression, and the *CaLDOX* gene expression was about three-fold lower in the *desi* seed coats and almost absent in the *kabuli* seed coats. *CaBAN* gene expression was negligible in the *kabuli* type seed coats (Fig. 1E; Fig. 2C). These results indicated that coloured flower petals of the *desi* type chickpeas possess a higher amount of anthocyanins and the seed coats contain a high level of proanthocyanidins in the inner integument and endothelial layers whereas, the petals and the seed coats of the *kabuli* type chickpeas did not show deposition of these flavonoid compounds. This observation is also supported by the expression of the genes indicative of anthocyanin and proanthocyanidin biosynthesis in the respective tissues of these two chickpea types.

**Figure 1.**
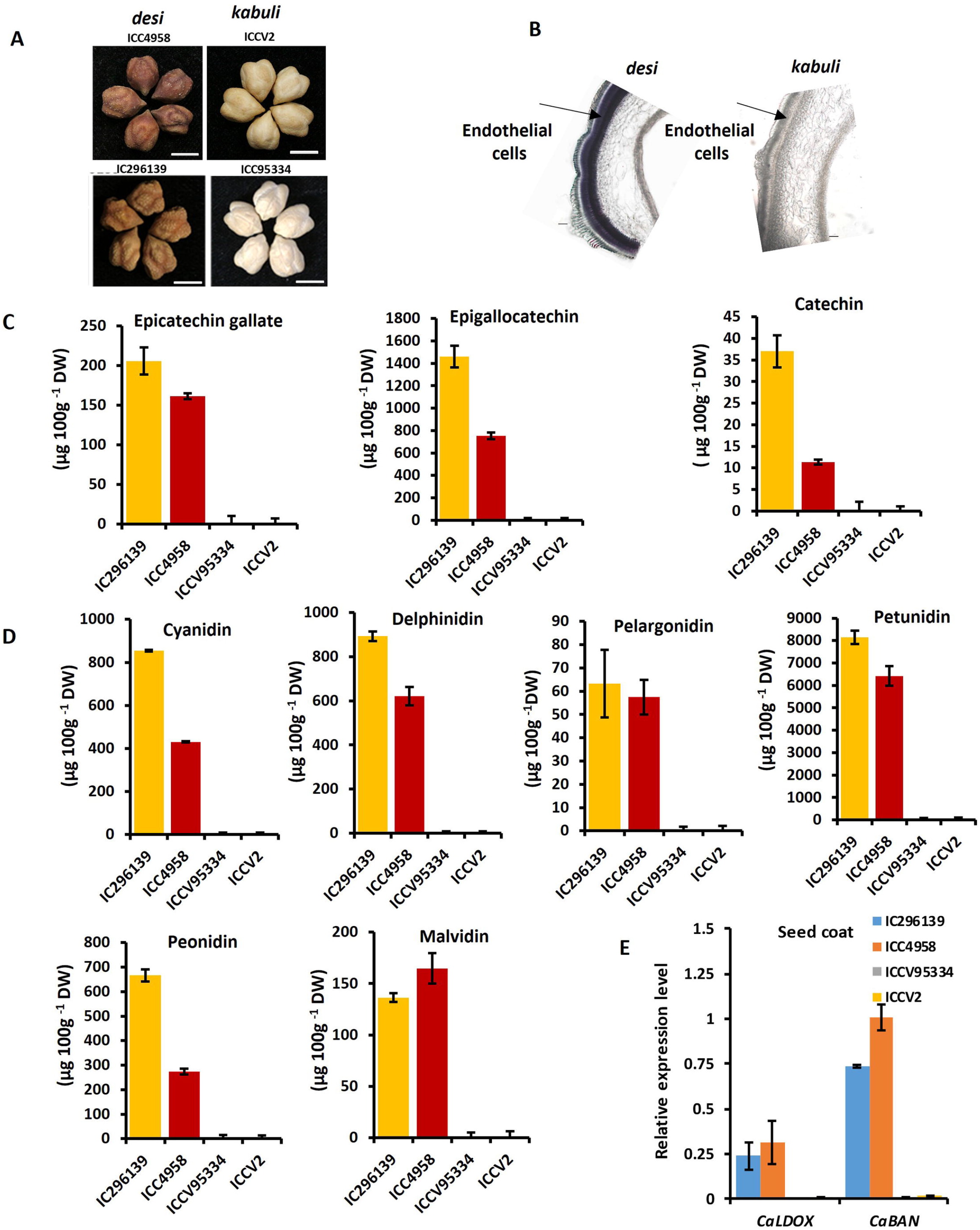
Phenotypic and biochemical studies of chickpea seed coat colour. **A**) Representative pictures of seeds of *desi* (accn: ICC4958, IC296139) and *kabuli* (accn: ICCV2, ICC95334). Scale bars, 5mm. **B**) Sections of *desi* and *kabuli* seed coats stained with 4-Dimethylaminocinnamaldehyde (DMACA) at 20x magnification, showing cellular structure including outer and inner testa. Scale bars, 100 µm. Quantification of **C**) Proanthocyanidin (PA) precursors epicatechin gallate, epigallocatechin and catechin and **D**) anthocyanin precursors cyanidin, delphinidin, pelargonidin1, petunidin, peonidin & malvidin in the seed coats of two *desi* (ICC4958 & IC296139) and *kabuli* (ICCV95334 & ICCV2) cultivars using LC-MS analysis. DW denotes dry weight. **E**) Relative expression of *CaLDOX and CaBAN* genes in seed coats of two *desi* and *kabuli* cultivars mentioned above as determined by qRT-PCR. *CaEF-1*α (NM_001365163.1) was used as internal control for qRT-PCR. All the assays were performed using three biological replicates, and the bars represent means ± SE.

**Figure 2.**
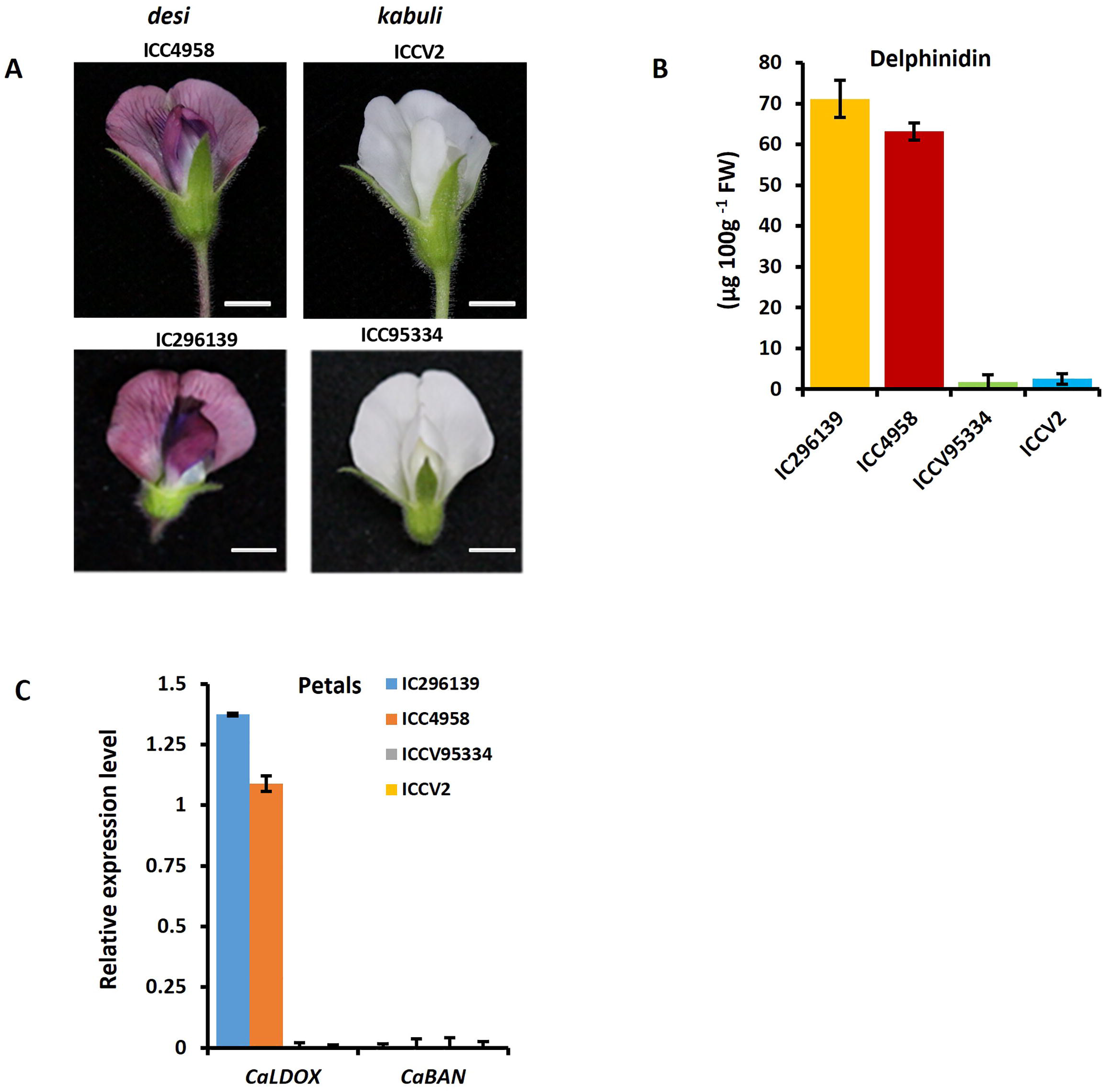
Phenotypic and biochemical studies of chickpea flower colour and relative expression analysis in desi and kabuli petal and seed coat. **A**) Representative pictures of flowers of *desi* (accn: ICC4958, IC296139) and *kabuli* (accn: ICCV2, ICC95334). Scale bars, 5mm. **B**) Estimation of delphinidinin the flower petals of *desi* (ICC4958 & IC296139) *kabuli* (ICCV95334 & ICCV2) cultivars. FW denotes fresh weight. **C**-**D**) Relative expression of *CaLDOX and CaBAN* genes in flower petals of two *desi* and *kabuli* cultivars mentioned above as determined by qRT-PCR. *CaEF-1*α (NM_001365163.1) was used as internal control for qRT-PCR. All the assays were performed using three biological replicates, and the bars represent means ± SE.

### Expression of the candidate *CabHLH* and *CaMATE1* genes

A gene (*CabHLH*, LOC101506726) encoding a sequence ortholog of *Medicago truncatula* TT8 (MtTT8) (Supplementary Fig. S2) was predicted to be associated with *kabuli* chickpea trait by GWAS (Penmetsa *et al*., 2016). Four coding sequences (CDS) variants (*b1* to *b4*) of this gene have been reported so far that are invariantly associated with the *kabuli* phenotype. Except for the *b2* allele, which is a missense mutation, all other alleles of the gene produce truncated proteins (Penmetsa *et al*., 2016). The *CabHLH* cDNAs of two *kabuli* accessions mentioned above were sequenced and it was found that ICC95334 possesses a 153 bp deletion causing premature termination and ICCV2 possesses a missense mutation replacing glutamine with proline (*b2* allele) (Supplimentary Fig. S3). The corresponding cDNA from the two *desi* accessions did not show any sequence variation.

The desi chickpea seed coat exhibited the highest expression of *CabHLH* gene followed by the mature flower petals as assessed by qRT-PCR (Fig. 3A). The leaves and the pod cover displayed a five-fold lower expression than the seed coat and the cotyledon, the lowest in the *desi* cultivar. The *kabuli* chickpea leaves and cotyledons showed a similar *CabHLH* gene expression to the *desi* chickpea while in the other tissues, expression of this gene was low. A promoter-reporter construct was made with 1717 bp upstream sequence of *CabHLH* gene (Supplementary Fig. S4) of ICC4958 (*desi*) fused to GUS (β-glucuronidase) and was used to develop transgenic chickpea lines. GUS-staining of different tissues of the stable T3 line corroborated the qRT-PCR result of tissue-specific gene expression patterns with the highest promoter activity in the seed coat (Fig. 3B). Subcellular localization of CabHLH protein fused to the green fluorescence protein (GFP) expectedly showed its localization in the nucleus (Fig. 3C).

**Figure 3.**
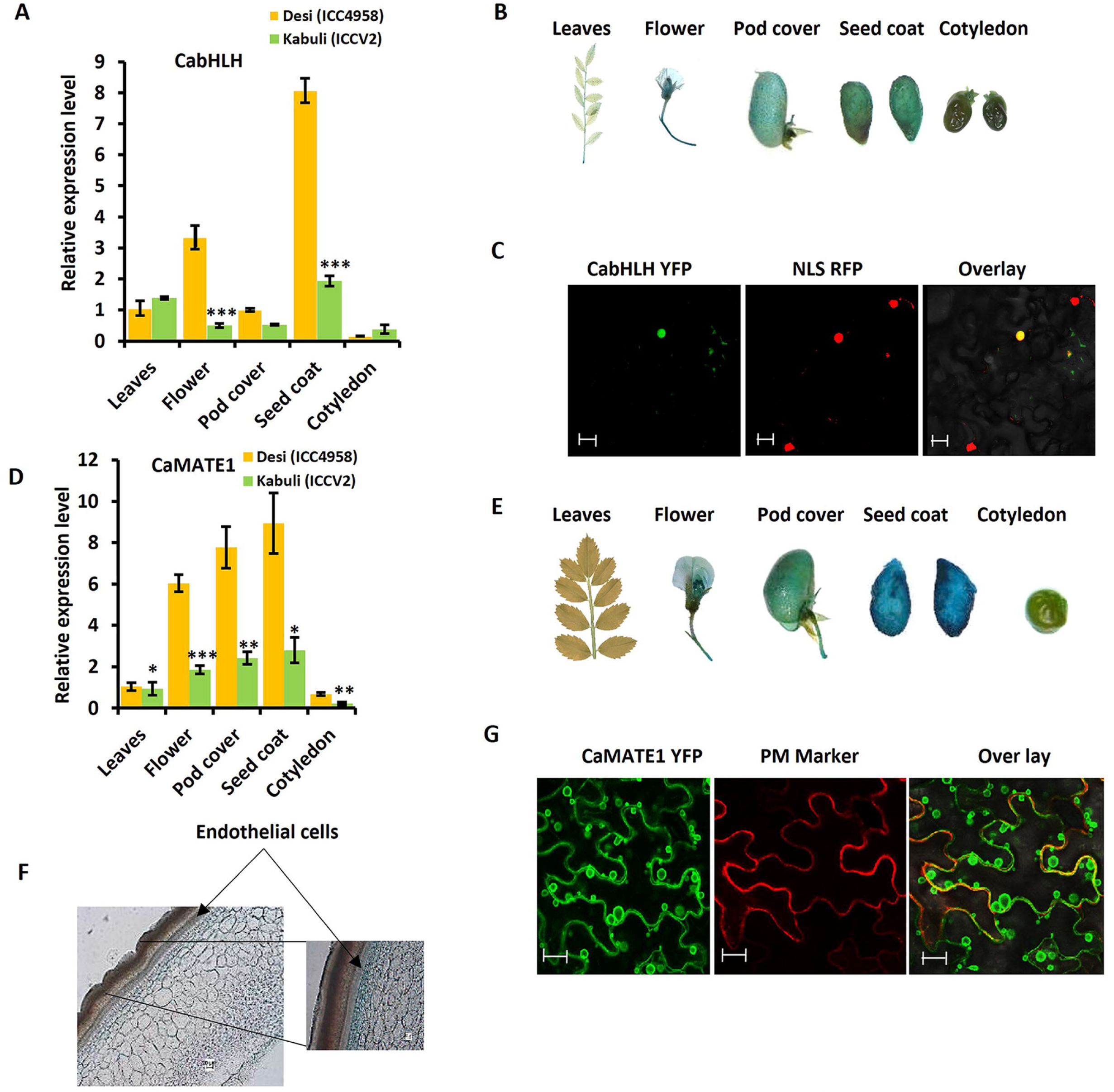
Relative expression of *CaMATE1* and *CabHLH* genes, their tissue-specific expression patterns and subcellular localization. **A**) qRT-PCR analysis of *CabHLH* gene expression in different tissues (leaves, flower, pod cover, seed coat and cotyledon) of *desi* (ICC4958) and *kabuli* (ICCV2) chickpea cultivar. *CaEF-1*α was used as an internal control and three biological replicates were used. **B**) *In planta* visualization of GUS (β*-glucuronidase*) activity in various tissues (leaves, flower, pod cover, seed coat and cotyledon) of T2 transgenic chickpea plants expressing *PromoterCabHLH::GUS* construct. **C**) Subcellular localization of CabHLH. CabHLH-YFP along with a nuclear marker (CD3-1106 mCherry) was transiently co-infiltrated in *N. benthamiana* leaves and was visualized under confocal microscopy after 60-65 hrs. Scale bars, 20 µm. **D**) qRT-PCR analysis of *CaMATE1*expression in different tissue of *desi* (ICC4958) and *kabuli* (ICCV2) chickpea cultivars. *CaEF-1*α was used as an internal control. Bars represent means ± SE. Asterisks indicate statistically significant differences between ICC4958 (*desi*) and ICCV2 (*kabuli*) determined using two-tailed Student’s *t*-test (* for *p* < 0.05, ** for *p* < 0.01, *** for *p* < 0.001). **E**) *In planta* visualization of GUS (β*-glucuronidase*) activity in various tissues (leaves, flower, pod cover, seed coat and cotyledon) of T2 transgenic chickpea plants expressing *PromoterCaMATE1::GUS* construct. **F**) Section of the seed coat of transgenic chickpea plants expressing GUS under *PromoterCaMATE1*. **G**) Subcellular localization of CaMATE1. *CaMATE1-YFP* along with plasma membrane (PM) marker (CD3-1007 mCherry) was co-agroinfiltrated in *N. benthamiana* leaves and was visualized under confocal microscope after 60-65 hrs. Scale bars, 20 µm. Co-localization with vacuolar membrane marker (CD3-975 mCherry) is presented in the supplementary figure S8B.

A genome-wide association study in chickpea has mapped three MATE encoding genes associated with seed coat colour (Bajaj *et al*., 2015). Two (LOC101503455, LOC101513987) of those are located on chromosome 2 and the other (LOC101501055) on chromosome 4. One of these proteins (encoded by LOC101513987) showed the highest sequence similarity with the MATE1 protein (MtTT12) of *M. truncatula*, (Supplimentary Fig. S5) which was reported as an essential membrane transporter for PA biosynthesis in the seed coat (Jhao and Dixon, 2009). This gene was used for further studies and is referred to as *CaMATE1*. The highest expression level of *CaMATE1* was observed in the seed coat followed by its expression in the pod cover and the flower petals of the *desi*-type plants. Its expression was very low in leaves and cotyledon in the qRT-PCR assay. The expression of *CaMATE* in *kabuli* type was approximately three-fold lower in seed coat, pod cover and flower petals in comparison to that in the desi type (Fig. 3D). Expression of the *GUS*-reporter gene under the control of a 1950 bp long upstream sequence of *CaMATE1* (Supplementary Fig. S6) gene in a stable transgenic T3 chickpea line followed by GUS-staining corroborated the qRT-PCR result supporting the expression of this gene mostly in the seed coat, pod cover, and flower with undetectable expression in leaves and cotyledons (Fig. 3E). *In planta* visulaization of GUS activity in the plants transformed with a GUS construct without any promoter did not show GUS-stain (Supplementary Fig. S7). Section of the GUS-stained seed coat showed *CaMATE1* promoter activity in the endothelial layer (Fig. 3F). However, the cross section of GUS-stained seed coat of CabHLH showed the stain in all the cells of seed (Supplimentary Fig. S8A). Expression of CaMATE1 protein fused to GFP indicated its localization in the tonoplast (Fig. 3G, Supplimentary Fig. S8B).

### Interaction of CabHLH with other proteins to form MBW complex

The transcription of the late biosynthetic genes governing anthocyanin and proanthocyanidin biosynthesis is regulated by MBW complex (Li *et al*., 2016). This ternary complex is composed of three transcription regulators belonging to R2R3-**M**YB, **b**HLH, and **W**D40 repeat families of proteins. Unlike bHLH and WD40 transcription factors, different MYB family members take part in regulating anthocyanin and PA biosynthesis in *Medicago*. LEGUME ANTHOCYANINE PRODUCTION 1 (LAP1) and PROANTHOCYANIN REGULATOR (PAR) are two MYB family TFs that function as the key regulators of anthocyanin and PA biosynthesis, respectively, in *M. truncatula* (Peel *et al*., 2009; Verdier *et al*., 2011). The orthologs of these MYB proteins were identified in chickpea and verified for their interactions using bimolecular fluorescence complementation (BiFC) in *Nicotiana benthamiana*. The CabHLH (XP_027189966.1, MtTT8 ortholog) and CaWD40-1 (XP_004502764.1, MtWD40-1 ortholog) of chickpea interacted with the chickpea MYB proteins CaLAP1 (XP_027188318.1, MtLAP1 ortholog) and CaPAR (XP_004507986.1, MtPAR ortholog) in the nucleus (Fig. 4A). However, the CabHLH protein of the *kabuli* chickpea accession ICCV2 carrying *b2* allele (CabHLH_*b2*) replacing a glutamine with proline at the 74^th^ position did not show any interaction with either CaLAP1 or CaPAR (Fig. 4B-C) suggesting Gln^74^ is essential for bHLH-MYB interaction to form MBW complex.

**Figure 4.**
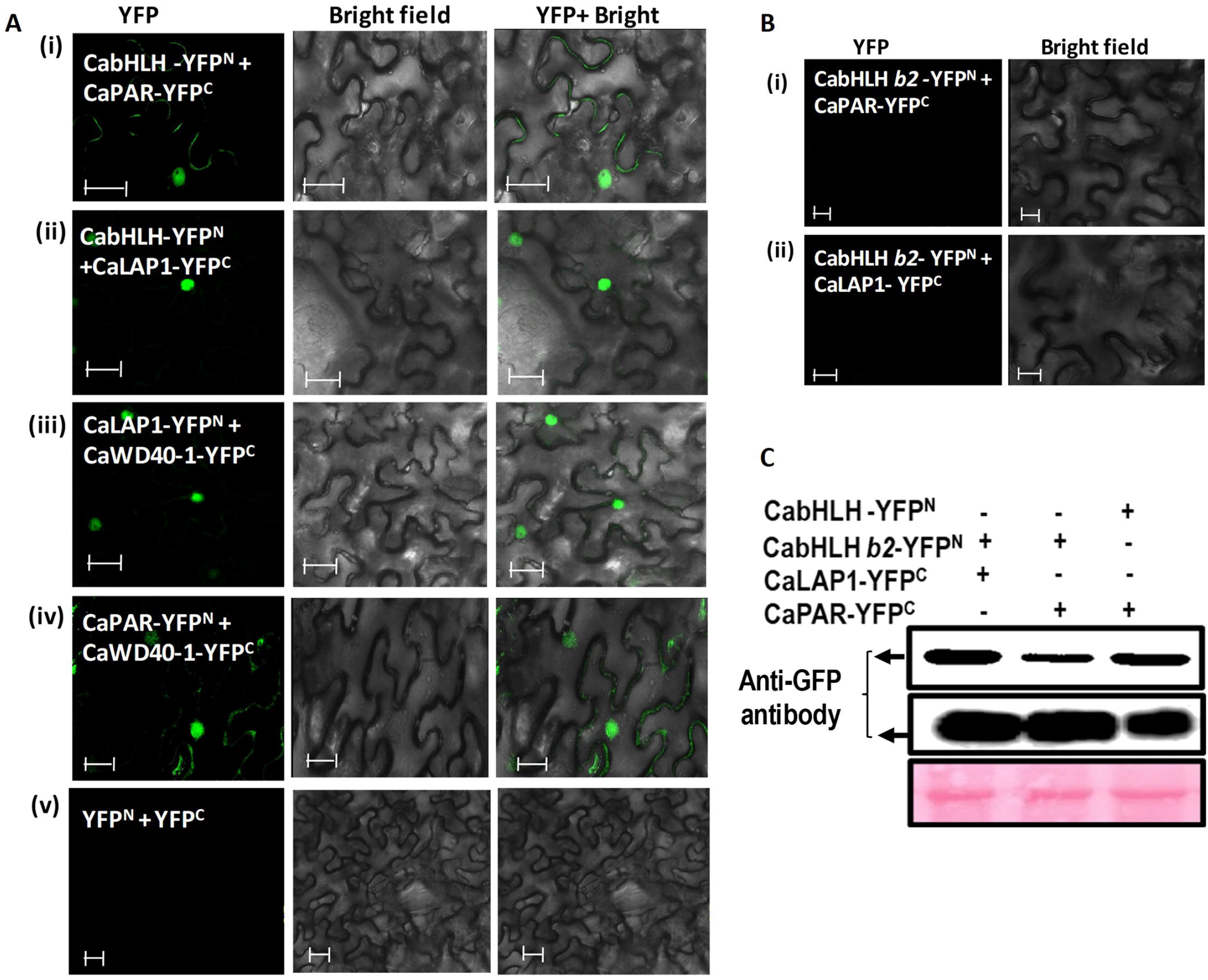
Biomolecular fluorescence complementation (BiFC) assay for physical interactions between CabHLH variants (CabHLH and CabHLH*b2*), CaMYB (CaPAR and CaLAP1) and CaWD40-1 proteins. **A**) The respective proteins were fused with either N- or C-terminal domain of yellow fluorescence protein (YFP) and expressed in combination as mentioned in the figure by co-infiltration in *N. benthamiana* leaves. **i-ii**) The interaction was shown between CabHLH together with CaPAR or CaLAP1. **iii-iv**) The interaction was shown between CaWD40-1 together with CaPAR or CaLAP1. **v**) The interaction was shown between the YFP^N^ (CD3-1651) and YFP^C^ (CD3-1648) empty vectors as negative control. **B**) CabHLH_*b2* when used for the interaction along with CaPAR or CaLAP1. The YFP represents the fluorescence field, the middle panel represents the bright field, and the rightmost panel represents the merged view of the fluorescent and bright field. **C**) Expression of proteins in the samples shown in B was demonstrated by immunoblotting with anti-GFP antibody in upper panel, lower panel represent the blotted membrane stained with Ponceau S. Co-infiltration of the constructs with their presence and absence was indicated with the sign + and – respectively. Scale bars, 20 µm.

### Assessment of transactivation potential of chickpea MBW complex

To assess the transactivation potential of the chickpea MBW complex and the specificity of CaLAP1 and CaPAR in activating the late anthocyanin- and PA-specific biosynthetic pathway genes, respectively, a co-transfection based transient expression system using protoplasts of the *Arabidopsis thaliana* At7 cell line was utilized. The normalized GUS reporter activity was taken as a measure of promoter activation. ORFs of *CabHLH, CaLAP1, CaPAR*, and *CaWD40-1* genes were fused with 35S CaMV promoter to use as effector constructs. The transactivation property of the effector constructs was tested with the reporter constructs harbouring of *CaDFR-2000* bp or *CaLDOX-1930* bp or *CaBAN-1816* bp promoters fused to the GUS reporter gene (Figure 5A). The transactivation assay indicated that CabHLH alone could not activate any of the test reporters. A combination of CabHLH, CaPAR, and CaWD40-1 significantly activated the promoter of *CaBAN* gene involved in PA biosynthesis. However, the replacement of CaPAR with CaLAP1 as an effector resulted in five-fold less activation of the *CaBAN* promoter indicating CaPAR is principally important for PA biosynthesis (Fig. 5B). Both the CaPAR and CaLAP1 in combination with CabHLH and CaWD40-1 were able to activate *CaLDOX* promoter, although the promoter activation with CaPAR was two-fold higher than that with CaLAP1 (Fig. 5C). On the other hand, MBW complex with CaLAP1 highly activated the promoter of Dihydroflavonol-4-reductase (*CaDFR*) gene, whereas the complex with CaPAR was ineffective on this promoter (Fig. 5D). Use of the *b2* allele of CabHLH (CabHLH*b2*, Glutamine^74^/Proline) in place the wild-type protein caused loss of transactivation activity of the MBW ternary complex indicating that the failure to form a functional MBW complex is the cause of the low anthocyanin and PA contents in the *kabuli* type chickpea varieties (Fig. 5B-D). Our experiment showed that CabHLH alone was unable to form active MBW complex to transactivate both anthocyanin and PA biosynthetic genes, however, separate MYB genes (CaLAP1 and CaPAR) were required for anthocyanin and PA biosynthesis.

**Figure 5.**
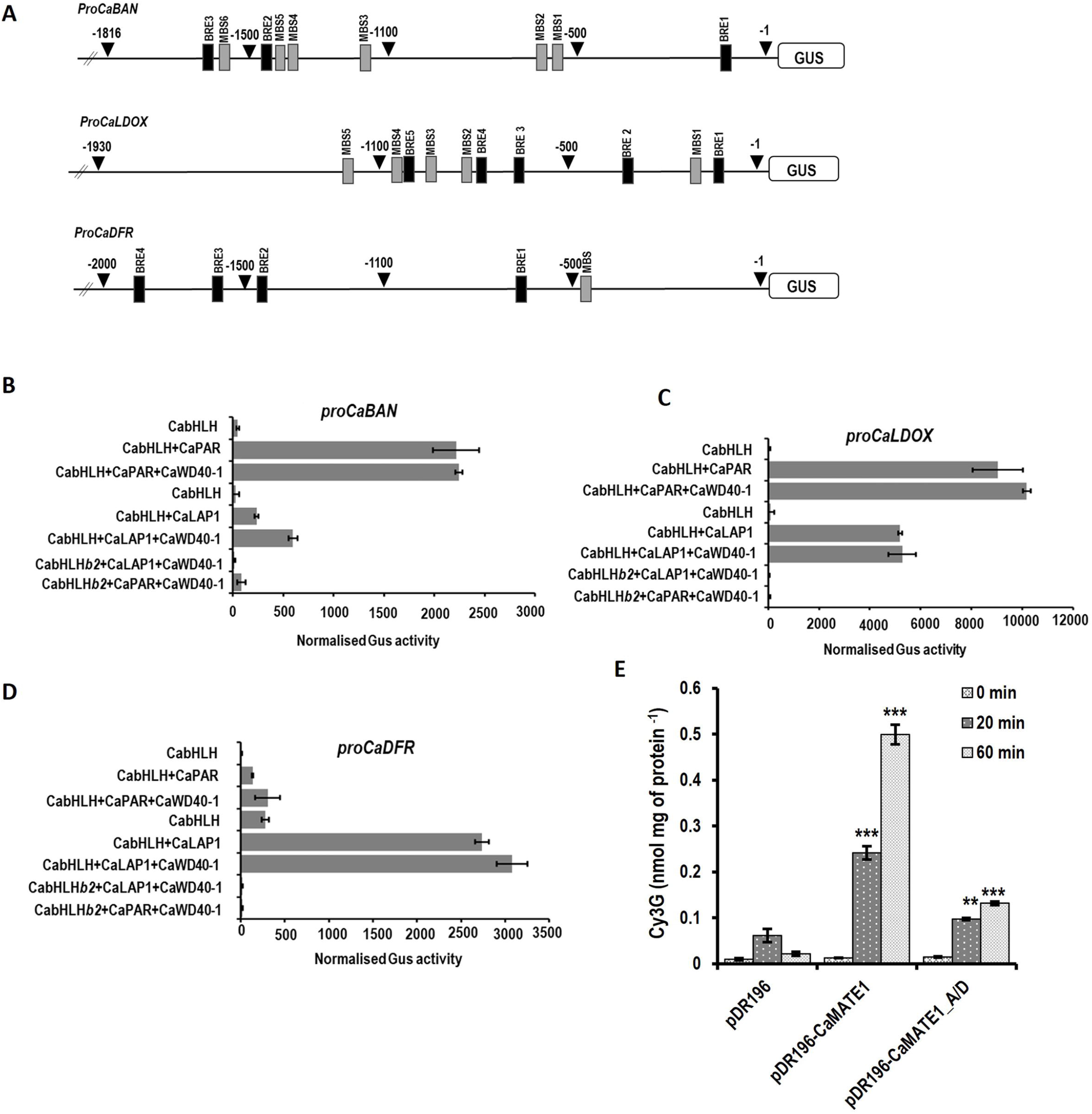
Transactivation assay of MBW transcription factor complex of CabHLH, CaMYB (CaPAR/CaLAP1) and CaWD40-1. **A**) Description of the GUS reporter constructs driven by the *CaLDOX* or *CaBAN* or *CaDFR* promoters used in the transactivation assay. bHLH and MYB responsive *cis*-acting elements such as, MYB binding sites (MBS) and bHLH responsive elements (BRE) were shown at positions relative to the transcription start sites. **B-D**) The effector constructs, CabHLH (CabHLH and CabHLH*b2*), CaPAR, CaLAP1 and CaWD40-1 were expressed under 35SCaMV promoter in the protoplasts of At7 cell lines along with the GUS reporter constructs in combinations as mentioned in the figures. Promoter activity was normalized with the luciferase activity of the coexpressed pUb14-2-Luciferase construct. Promoter activities of the reporter constructs were presented in the form of normalized GUS activity. All the data were obtained from three biological replicates and expressed as means ± SE. Asterisks indicate statistically significance by two-tailed Student’s *t*-test (*** for *p* < 0.001). Assessment of uptake activity of cyanidin 3’O-glucoside (Cy3G) by yeast membrane vesicle cells. **E**) Measurement of uptake of Cy3G into the membrane vesicles derived from *S. cerevisiae* AD12345678 cells expressing the empty vector pDR196 (EV), CaMATE1 and CaMATE1^A/D^ at 0 min, 20 min and 60 min after addition of the substrate. Results are mean and ± SE from three biological replicates. Asterisks indicate statistically significance by two-tailed Student’s *t*-test (** for *p* < 0.01, *** for *p* < 0.001).

### Assessment of transport potential of the natural alleles of CaMATE1

Loss-of-function mutation of *Arabidopsis* TT12, the tonoplastic MATE transporter, produced transparent testa phenotype. When expressed in yeast, *Arabidopsis* TT12 facilitated transport of cyanidin 3-*O*-glucoside (Cy3G) in the membrane vesicles, like its ortholog in *M. truncatula* (MtMATE1) (Zhao and Dixon, 2009). To assess the capability of tonoplast localized CaMATE1 to transport Cy3G, *CaMATE1* was expressed in the *Saccharomyces cerevisiae* strain AD12345678 lacking multidrug resistance activity (Morita *et al*., 2009). Transport assay with the microsomal fraction showed CaMATE1 was able to transport Cy3G through the vacuolar membrane (Fig. 5E). A genome-wide association study predicted a strong association potential of one non-synonymous SNP (C/A) in CaMATE1 with the light seed coat colour (Bajaj *et al*., 2015). This SNP substituted alanine (A) in *desi* chickpeas with aspartic acid (D) in the *kabuli* chickpeas at the 117^th^ position in the conserved transport domain of CaMATE1 (Supplementary Fig. S9). The yeast microsomal fraction expressing CaMATE1^A/D^ carrying aspartic acid at 117^th^ position exhibited 3.7-fold lower vacuolar uptake activity (Fig. 5E), explaining the association of a natural allele of CaMATE1 with the seed coat colour.

### Role of CabHLH and CaMATE1 in determining flower and seed coat colour

In order to investigate the role of CabHLH in determining flower and seed coat colour in the native system chickpea, the expression of this gene was silenced by RNA interference (RNAi) (Supplementary Fig. S10A). The RNAi construct was developed with 200 bp of coding sequence *CabHLH* (Supplementary Fig. S10B; Supplementary Table S1) under the control of *CaMATE1* promoter as this promoter was active only in flower petals, pod cover, and seed coat. The construct was used to transform a *desi* type chickpea accession ICC4958 and four independent T4 homozygous lines were taken for further studies. A reduction of 60%-80% in the *CabHLH* expression in the petals and seed coat was observed in all the lines (Fig. 6A; Fig. 7A). Subsequently, expression levels of *CaLDOX* and *CaBAN* genes in these lines were more than 2-5-fold lower in the petals and seed coats (Figure 6A). Moreover, the relative expression analysis of *CaDFR* gene in petal and seed coat showed >10 fold and 3-4-fold change respectively in *CabHLH* RNAi lines (Fig. 6A; Fig. 7A). These lines produced pale-brown coloured seeds with a smooth texture like the kabuli seeds however, those seeds were angular in shape like the *desi* seeds (Fig. 6B). These seed coats showed a 3-5-fold lower deposition of insoluble and soluble DMACA-reactive compounds in the endothelial layer as compared to their parent accession ICC4958 (Fig. 6C-G). The gene silenced chickpea seeds also showed reduced mucilage accumulation (Fig. 6E). The contents of PA precursors, catechin, epigallocatechin and epicatechingallate; and the anthocyanin precursors, cyanidin, delphinidin, pelargonidin, petunidin and malvidin were reduced by 5-20-fold in the seeds of RNA-silenced lines (Fig. 6H-I). The flower petals of the RNAi lines were almost white except for the light purple colour in the veins and the tip of the inner whorl (Fig. 7B). Delphinidin content was reduced by 3-6-fold in these lines (Fig. 7C). Total anthocyanin content (TAC) was also reduced 3-5-fold in seed coat and petals of these lines (Supplementary Fig. S11A-B). There were no other apparent phenotypic changes in the vegetative or reproductive characteristics of the transgenic lines as compared to their parent accession ICC4958 (Supplementary Fig. S12).

**Figure 6.**
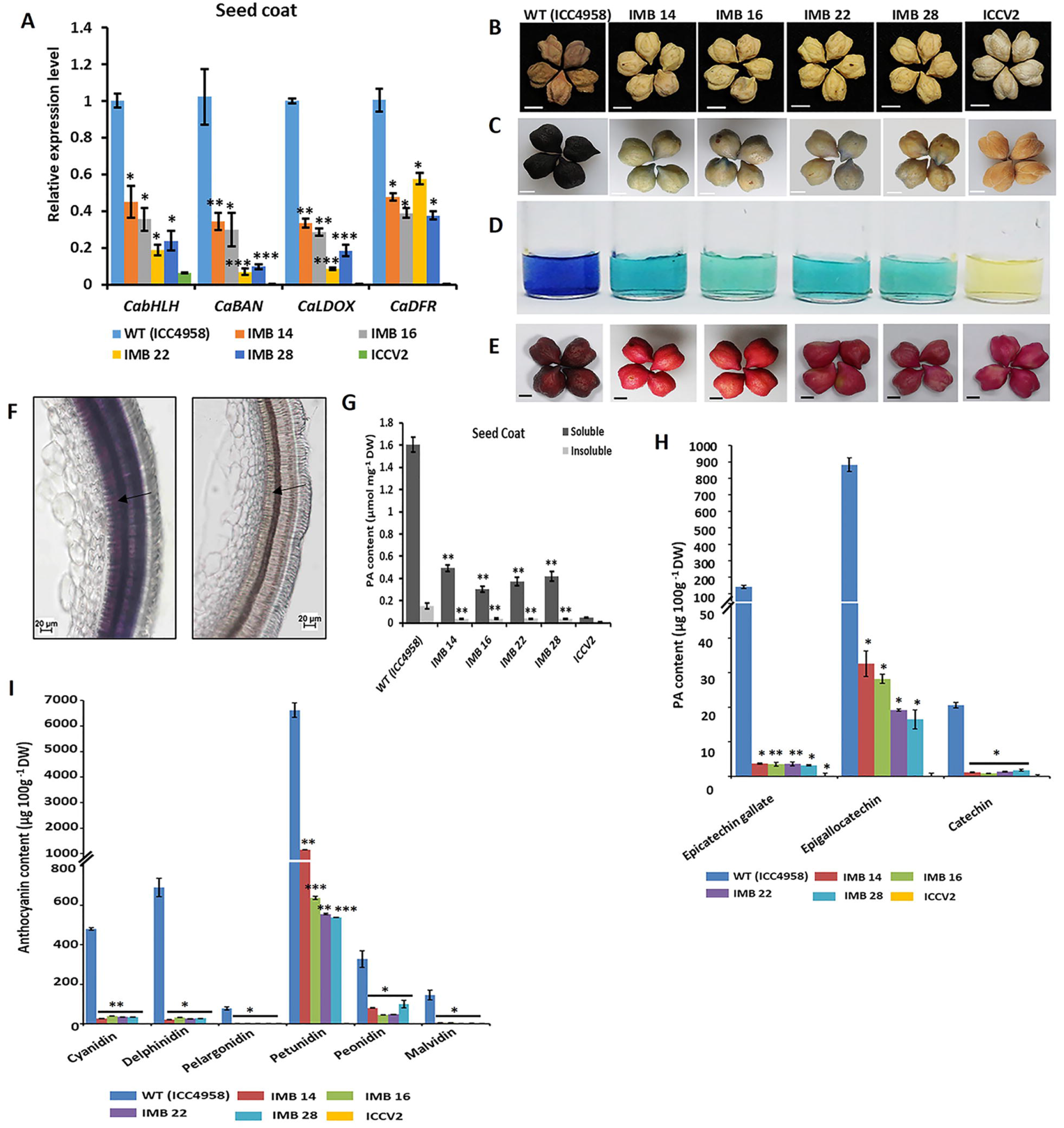
Knockdown of *CabHLH* gene expression and quantitative analysis of anthocyanin and proanthocyanidin in seed coat of chickpea accession ICC4958 (*desi*) by RNA interference (RNAi) and their characterization in the stable chickpea transgenic lines. **A**) Expression analysis of *CabHLH* gene and its downstream targets *CaBAN, CaLDOX*, and *CaDFR* in four independent transgenic lines, WT (ICC4958) and ICCV2 (*kabuli*) cultivars in seed coats. *CaEF-1*α was used as the internal control. **B**) Seed phenotype of four independent T4 lines (IMB14, IMB16, IMB22 & IMB 28). Scale bars, 5 mm. **C**-**D**) Staining of whole seeds and estimation of total soluble PA in the seed coats by DMACA. Scale bars, 5 mm. **E**) Ruthenium red staining of whole seeds to detect mucilage accumulation in four RNAi lines. **F**) DMACA-stained sections of the seed coats of chickpea accession ICC4958 (WT) (left) and the IMB16 line (right) at 20x magnification to show the accumulation of PAs in the endothelium layer (Arrows). Scale bars, 20 μm. **G**) Estimation of total soluble and insoluble PAs in the seed coats of RNAi lines. Estimation of **H**) PA precursors (epicatechin gallate, epigallocatechin and catechin) and **I**) anthocyanin precursors (cyanidin, delphinidin, petunidin, peonidin, malvidin) in the seed coats of the chickpea WT (ICC4958) and the RNAi lines by LC-MS analysis. DW denotes dry weight. Bars in represent means ± SE. Asterisks indicate statistically significant differences from chickpea accession ICC4958 (WT) determined using two-tailed Student’s *t*-test (* for *p* < 0.05, ** for *p* < 0.01, *** for *p* < 0.001).

**Figure 7.**
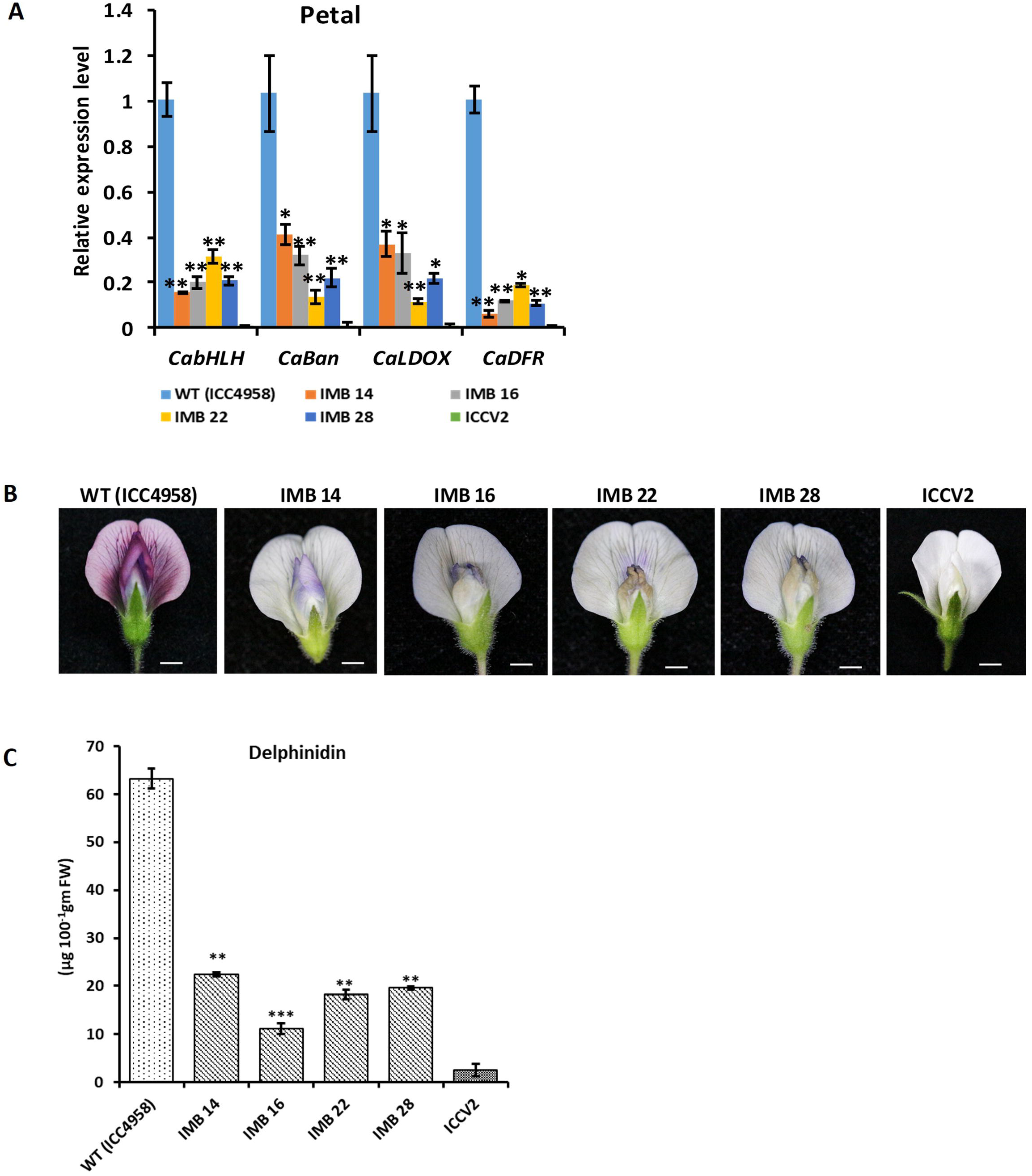
Knockdown of *CabHLH* gene expression and quantitative analysis of anthocyanin and proanthocyanidin in flower petals of chickpea accession ICC4958 (*desi*) by RNA interference (RNAi) and their characterization in the stable chickpea transgenic lines. **A**) Expression analysis of *CabHLH* gene and its downstream targets *CaBAN, CaLDOX*, and *CaDFR* in four independent T4 transgenic lines (IMB14, IMB16, IMB22 & IMB 28)), WT (ICC4958) and ICCV2 (*kabuli*) cultivars in flower. *CaEF-1*α was used as the internal control. **B**) Phenotype of flowers of chickpea WT (ICC4958), ICCV2 as control and four *CabHLH* RNAi lines. A representative image of each line has been presented. Scale bars, 5 mm. **C**) Quantitative estimation of delphinidin in the flower petals of the lines mentioned above. FW denotes fresh weight. All data are from three biological replicates and represent means ± SE. Asterisks indicate statistically significant differences from the chickpea WT (ICC4958) determined using two-tailed Student’s *t*-test (* for *p* < 0.05, ** for *p* < 0.01, *** for *p* < 0.001).

Similarly, four independent RNAi lines were developed for *CaMATE1* using 200 bp regions of its cDNA (Supplementary Fig. S10C; Supplementary Table S1). Tissue-specific expression analysis showed down-regulation of *CaMATE1* gene by 4-5-fold in flower and seed coat (Fig. 8A; Fig. 9A). No downreguation of others *CaMATE* genes (LOC101503455, LOC101501055) was seen in the flower petal and seed coat of the *CaMATE1*-downregulated lines (Supplementary Fig. S13A-B). The T5 homozygous lines showed similar flower and seed phenotypes as we observed in the case of *CabHLH* RNAi transgenic lines with no compromise in vegetative or reproductive characteristics (Fig. 8B, Fig. 9B; Supplementary Fig. S14). Retention of light purple colour in the petal veins and at the tip of the second whorl in the transgenic lines in both the gene-silenced chickpea lines was probably due to the residual leaky expression of the genes concerned. Total soluble and insoluble PA estimation result revealed 3-4-fold less accumulation of PA in the endothelial layer of the seed coat of transgenic lines by DMACA mediated estimation and staining (Fig. 8C-F). The seed coat of these lines exhibited very low levels of PA and anthocyanin contents (Fig. 8G-H; Supplementary Fig. S11C). The flower was almost white as observed in the *CabHLH* RNAi lines and these lines exhibited 4-5-fold reduced delphinidin content and reduced total anthocyanin content (TAC) in petals. (Fig. 9B-C; Supplementary Fig. S11D). The results described above provided the molecular evidence that *CabHLH* (*MtTT8*) and *CaMATE1* (*MtMATE1*) are essential for determining both the flower and seed coat colours of chickpea and tissue-specific independent genetic manipulation of any of these two genes can alter the flower and seed coat colours of chickpea.

**Figure 8.**
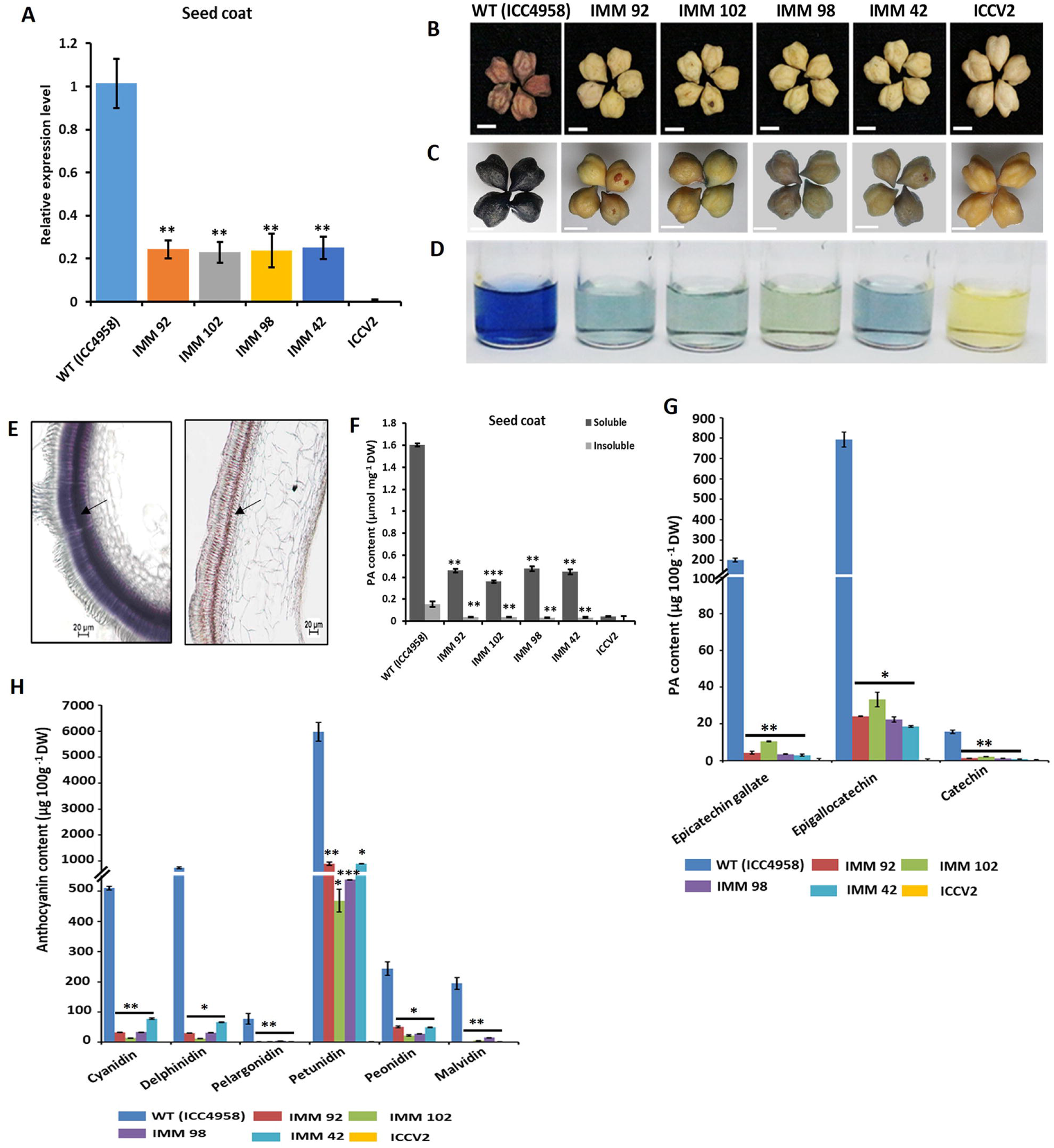
Knockdown of *CaMATE1* gene expression and quantitative analysis of anthocyanin and proanthocyanidin in chickpea accession ICC4958 (*desi*) by RNA interference (RNAi) and their characterization in the stable chickpea transgenic lines. **A**) Expression analysis of *CaMATE1* gene in four independent RNAi lines, WT (ICC4958) and ICCV2 (*kabuli*) cultivars in seed coats. *CaEF-1*α was used as the internal control. **B**) Seed phenotype of four independent T4 lines (IMM42, IMM92, IMM98 and IMM102). Scale bars, 5 mm. **C**-**D**) Staining of whole seeds and estimation of total soluble PA in the seed coats by DMACA in four RNAi lines. Scale bars, 5 mm. **E**) DMACA-stained sections of the seed coats of chickpea accession ICC4958 (WT) (left) and the IMM92 line (right) at 20x magnification to show the accumulation of PAs in the endothelium layer (Arrows). Scale bars, 20 μm. **F**) Estimation of total soluble and insoluble PAs in the seed coats of RNAi lines. Estimation of **G**) PA precursors (epicatechin gallate, epigallocatechin and catechin) and **H**) anthocyanin precursors (cyanidin, delphinidin, petunidin, peonidin, malvidin) in the seed coats of the chickpea WT (ICC4958) and the RNAi lines by LC-MS. DW denotes dry weight. Bars in represent means ± SE. Asterisks indicate statistically significant differences from chickpea accession ICC4958 (WT) determined using two-tailed Student’s *t*-test (* for *p* < 0.05, ** for *p* < 0.01, *** for *p* < 0.001).

**Figure 9.**
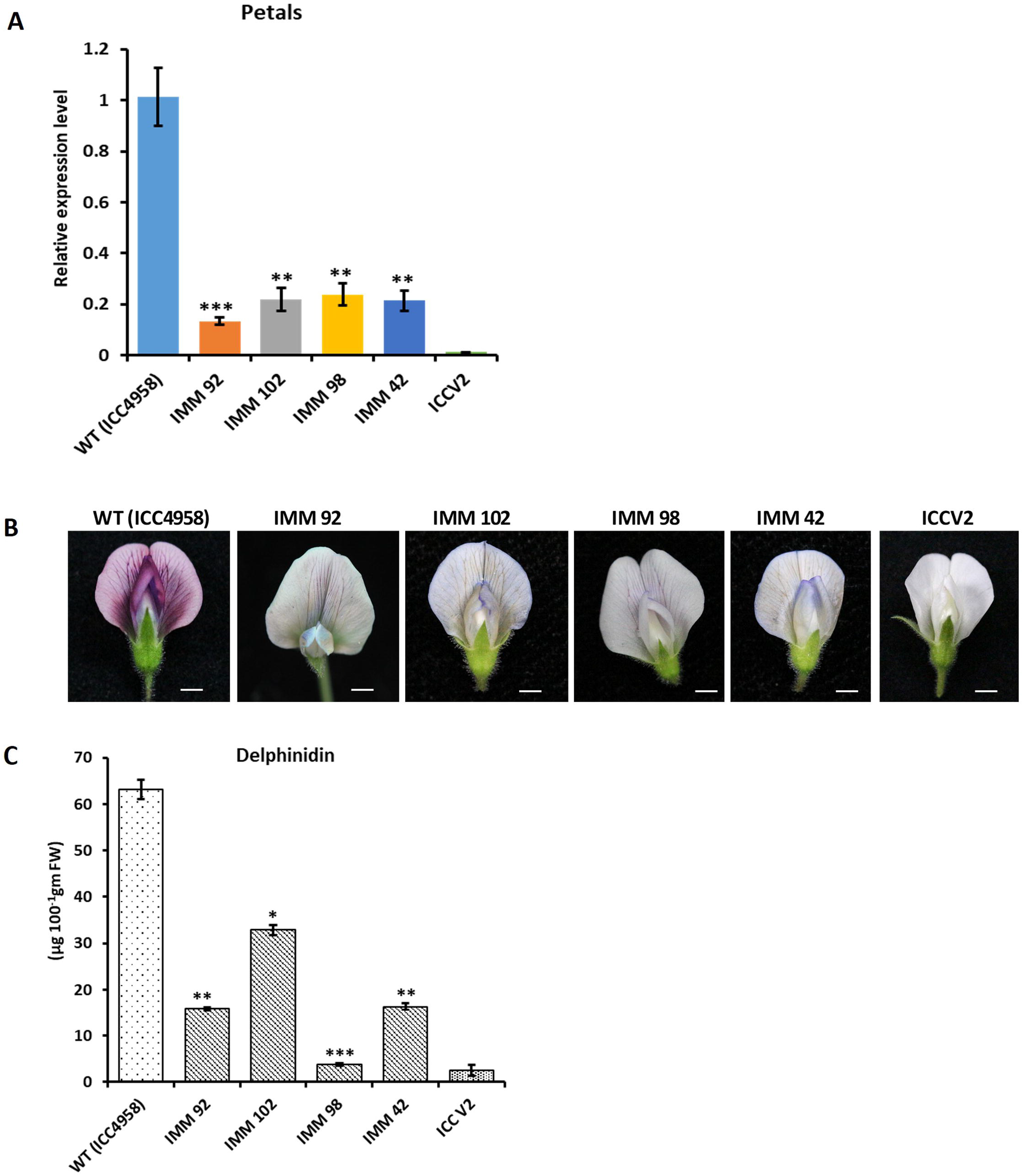
Knockdown of *CaMATE1* gene expression and quantitative analysis of anthocyanin and proanthocyanidin in flower petals of chickpea accession ICC4958 (*desi*) by RNA interference (RNAi) and their characterization in the stable chickpea transgenic lines. **A**) Expression analysis of *CaMATE1* gene in flower petals of four T4 independent RNAi lines (IMM42, IMM92, IMM98 and IMM102), WT (ICC4958) and ICCV2 (*kabuli*) cultivars. *CaEF-1*α was used as the internal control. **B**) Phenotype of flowers of chickpea WT (ICC4958), ICCV2 as control and four *CaMATE1* RNAi lines. A representative image of each line has been presented. Scale bars, 5 mm. **C**) Quantitative estimation of delphinidin in the flower petals of the lines mentioned above. FW denotes fresh weight. All data are from three biological replicates and represent means ± SE. Asterisks indicate statistically significant differences from the chickpea WT (ICC4958) determined using two-tailed Student’s *t*-test (* for *p* < 0.05; ** for *p* < 0.01, *** for *p* < 0.001).

## Discussion

The study described above provided some novel information in addition to linking the chickpea flower- and seed coat colours by two individual genes of a metabolic pathway. The role of bHLH family genes in anthocyanin and PA biosynthesis has been demonstrated in other plants. For example, in *Arabidopsis*, EGL3 (bHLH) and PAP1 (MYB) interact to form MBW complex with TTG1 to control the anthocyanin biosynthesis while, TT8 (bHLH) and TT2 (MYB) and TTG1 control the PA biosynthesis. We have shown that a single gene CabHLH was capable of regulating both the PA and anthocyanin biosynthesis in chickpea and this chickpea ortholog of MtTT8 regulate both the colours of seed coats and flower. We have shown using co-transfection assay that a single bHLH protein (CabHLH) interacts with two different types of MYBs (LAP1 and PAR) to form a functional MBW complex and regulates anthocyanin and PA biosynthetic genes, respectively. Similarly, the role of MATE1 (TT12) in the regulation of flower colour has not been demonstrated in *Arabidopsis* and *Medicago*. We have shown, CabHLH and CaMATE1 control two visible and agronomically important traits, flower colour and seed coat colour in chickpeas. The natural alleles of *CabHLH* and *CaMATE1* that we studied here are not available in their orthologs in other studied species. Previously, these chickpea genes were identified as polymorphic and those GWAS studies indicated the association of these genes and the alleles with these traits, however, using various different genetic backgrounds. In the present study, based on molecular and biochemical studies, we provided experimental evidence explaining how these natural alleles causing single amino acid replacements impaired these genes’ functions and changed the flower and seed phenotypes.

Flower and seed colours are the oldest traits used in the study of genetics and heredity. Contrasting external appearance and simple inheritance are the reasons for the choice. The same traits also formed one of the bases for the selection and domestication of chickpea. Condensed tannin in the seed coat causes roughness in the texture and adds astringency in taste (Soares *et al*., 2020). This could be another major reason for preference and domestication of smooth and light brown seeded *kabuli* chickpeas. White-flowered genotypes are not reported so far in the primary (*C*.*reticulatum*) and secondary (*C*.*echinospermum*) gene pools of chickpea. It is known only in the perennial *Cicer* species in the tertiary gene pool (Van Oss *et al*., 2015). Therefore, natural mutation at multiple positions of *CabHLH* gene and independent selection of those alleles at different geographic locations have been proposed for the origin of *kabuli* genotypes. Although, it is thought that the kabuli phenotype emerged from the desi chickpea during domestication (Penmetsa *et al*., 2016), Toker (2009) has shown by radiation-mediated mutation that kabuli phenotype can be created directly from the wild progenitor (*C. reticulatum*) of cultivated chickpea suggesting an additional and/or alternative path for evolution of kabuli type by spontaneous mutation of the wild progenitor. The other general growth pattern-related differences between the *desi* and the *kabuli* genotypes are probably due to the varietal and agro-climatic differences.

Annotation of the present chickpea genome assemblies (NCBI Bioproject ID: PRJNA175619, PRJNA78951) shows the presence of twenty-five genes encoding MATE proteins in chickpea. A tandem cluster of *MATE* genes (*TT12* sequence orthologs) occurs in a narrow interval of chromosome 4 showing high purification selection within the *desi* cultivars (Varshney *et al*., 2013). Many MATE transporters localize in the vacuole membrane and transport secondary metabolites including alkaloids, and multiple MATE transporters might be involved in the transport of PA and anthocyanin precursors (Upadhyay *et al*., 2019). We selected one of the three *MATE* genes which were previously shown to be associated with seed colour for our work due to its sequence homology with the *M. truncatula* MATE1, which functionally complemented a *Arabidopsis tt12* mutant (Jhao and Dixon, 2009; Bajaj *et al*., 2015). Interestingly, the CabHLH-silenced and CaMATE1-silenced chickpea lines showed similar reduced PA profiles in the seed coat. It is in contrast to *M. truncatula* in which genetic loss of MtMATE2, which is predominantly expressed in leaf and flower, caused a reduction of anthocyanin pigmentation only in leaf and flower due to a decrease in various flavonoids, however, led to increased PA biosynthesis in seeds (Zhao *et al*., 2011). It appears that as described in the case of MtMATE1, CaMATE1 is an essential membrane transporter for the precursors of PA and anthocyanin both in chickpea flower petals and seed coat. Its tissue-specific expression pattern also suggests this. We have assessed the ability of CaMATE1 to transport an anthocyanin derivative and shown a compromised activity of a natural allele of CaMATE1, which was associated with light seed coat colour. The other MATE transporters are probably important in other tissues or for other functions.

We have functionally characterized the important genes involved in PA and anthocyanin biosynthesis in chickpea flower and seed coat. Decreased *CabHLH* expression in flower petal and seed coat resulted in reduced expression of *CaBAN* and *CaLDOX*. This showed that CabHLH transcriptionally regulates a part of the flavonoid pathway that produces PA and anthocyanin. As shown in *M. truncatula*, CabHLH and CaWD40-1, along with two MYB family transcription factors, CaPAR and CaLAP1 form the active ternary MBW transcription complex for activation of the promoters of the late PA and anthocyanin biosynthetic genes *CaBAN, CaLDOX* and *CaDFR*. Besides this, silencing of *CabHLH* expression reduced seed coat mucilage. Although MtTT8 is also involved in the regulation of foliar anthocyanin (Pang *et al*., 2009; Li *et al*., 2016), we did not observe this phenotype in the gene silenced lines probably due to tissue-specific gene silencing. The trans-activation experiment results suggested, CabHLH together with CaLAP1 or CaPAR and CaWD40-1 could regulate the expression of the *CaLDOX* promoter. However, CaPAR was found to specifically regulate *CaBAN* promoter while CaLAP1, specifically regulated *CaDFR* promoter in our experiment. The natural allele of CabHLH (*b2*, Gln/Pro) was not able to activate these promoters suggesting that the formation of MBW complex was essential and the Gln^74^ was essential to form the complex between CabHLH and the MYB proteins CaLAP1 and CaPAR. Overall, our results suggested that the basic molecular mechanism of PA and anthocyanin biosynthesis as described in the model legume *M. truncatula* is more or less conserved in chickpea. However, chickpea provided a unique opportunity of studying flower and seed coat colours together as these traits are genetically linked. Our results also showed that these same genes are equally critical in the biosynthesis of PA and anthocyanin which are responsible for flower colour in chickpea. Although the seed colour of the gene-silenced lines is similar to those of the *kabuli* chickpeas, the seeds were angular in shape like those of the non-transformed desi accession suggesting seed shape is regulated by other genes. However, the low-PA gene-silenced lines produced seeds with a smooth texture like the *kabuli* type which indicates the roughness of the desi chickpea seeds is due to high tannin content. We did not observe any apparent variation in the reproductive phenotypes of the gene silenced lines such as changes in size or shape or number of flowers and seeds. It appears that these genes are not critical for the reproductive behaviour of chickpea. There are possibilities that these genes perform additional functions in other tissues and genetic disruption of these two genes may lead to phenotypic variations in the vegetative parts of the plant.

Taken together, our results provide molecular evidence that flower and seed coat colours, the most commercially important agronomic traits of a highly important legume crop chickpea are biochemically regulated by the same genes and these traits can be altered by manipulation of a single gene.

## Supplementary data

Supplementary Fig. S1. Relative expression analysis of *CaActin* with *CaEF-1*α.

Supplementary Fig. S2. Phylogenetic tree of CabHLH and its homologs.

Supplementary Fig. S3. Clustal-W gene alignment of *bHLH* and *b4* (ICC95334) and *b2* (ICCV2) allele.

Supplementary Fig. S4. Promoter sequence of *CabHLH* used for *In-planta* expression study.

Supplementary Fig. S5. Phylogenetic tree of CaMATE and its homologs.

Supplementary Fig. S6. Promoter sequence of *CaMATE1* used for *In-planta* expression study.

Supplementary Fig. S7. *In planta* visualization of without promoter GUS activity.

Supplementary Fig. S8. Cross-section of *PromoterCabHLH::GUS* construct transgenic seed lines and subcellular localization of CaMATE1 protein.

Supplementary Fig. S9. Sequence alignment of MATE1 from different plant species.

Supplementary Fig. S10. Schematic representation of RNAi cassette.

Supplementary Fig. S11. Total anthocyanin estimation (TAC) in flower and seed coat of both RNAi lines.

Supplementary Fig. S12. Phenotypic evaluation of *PromoterMATE1::CabHLH* RNAi transgenic lines.

Supplementary Fig. S13. Relative expression analysis of other two *CaMATE* genes.

Supplementary Fig. S14. Phenotypic evaluation of *PromoterMATE1::CaMATE1* RNAi transgenic lines.

Supplementary Table S1. List of primers used in this study.

## Acknowledgements

The assistance of central instrumentation facilities, metabolomics facility and DISC of NIPGR in various experiments is acknowledged. The authors are grateful to the DBT-eLibrary Consortium (DeLCON) for providing access to e-Resources.

## Authors contributions

DC initiated, conceived, designed, and coordinated the research project and wrote the manuscript. AP designed the metabolomics, transactivation and transport assay experiments. LP has performed and analysed all the data. VD and SKG contributed in generating chickpea transgenic lines and phenotypic evaluation. SS has contributed in LC-MS analysis.

## Conflict of interest

Authors declare no conflict of interest. All authors read and approved the final manuscript.

## Funding Statement

The project is funded by the National Institute of Plant Genome Research, Department of Biotechnology (DBT), Ministry of Science and Technology, Government of India, J.C. Bose Fellowship (JCB/2020/000014) from Science and Engineering Research Board, Department of Science and Technology. LP, & SS are supported by fellowships from Council of Scientific and Industrial Research (CSIR, Govt. of India) and DBT, respectively. The funders had no role in study design, data collection and analysis, decision to publish, or preparation of the manuscript.

## Data availability

All data supporting the findings of this study are available within the paper and within its supplementary materials published online. The gene sequences are available in National Center for Biotechnology Information (NCBI) (http://www.ncbi.nlm.nih.gov).

## Abbreviations

BAN: BANYULS
BiFC: Bimolecular fluorescence complementation
bHLH: basic helix-loop-helix
Ca: *Cicer arietinum*
CDS: Coding sequence
Cy3G: Cyanidin 3-O-glucoside
DMACA: Dimethylaminocinnamaldehyde
DFR: Dihydroflavonol-4-reductase
GFP: Green fluorescent protein
LAP1: Legume Anthocyanine Production 1
LC-MS: Liquid chromatography-coupled-mass spectrometry
LDOX: Leucoanthocyanin Dioxygenase
MATE: Multidrug and toxic compound extrusion
Mt: Medicago truncatula
MBW complex: MYB- bHLH- WD40 comlex
PAs: Proanthocyanidins
PAR: Proanthocyanin Regulator
qRT-PCR: Quantitative Real time PCR
TAC: Total anthocyanin content.

## Notes

### Competing Interest Statement

The authors have declared no competing interest.

