## Supplementary Information for "Biochemical analysis of anthocyanin and proanthocyanidin and their regulation in determining chickpea flower and seed coat colours"

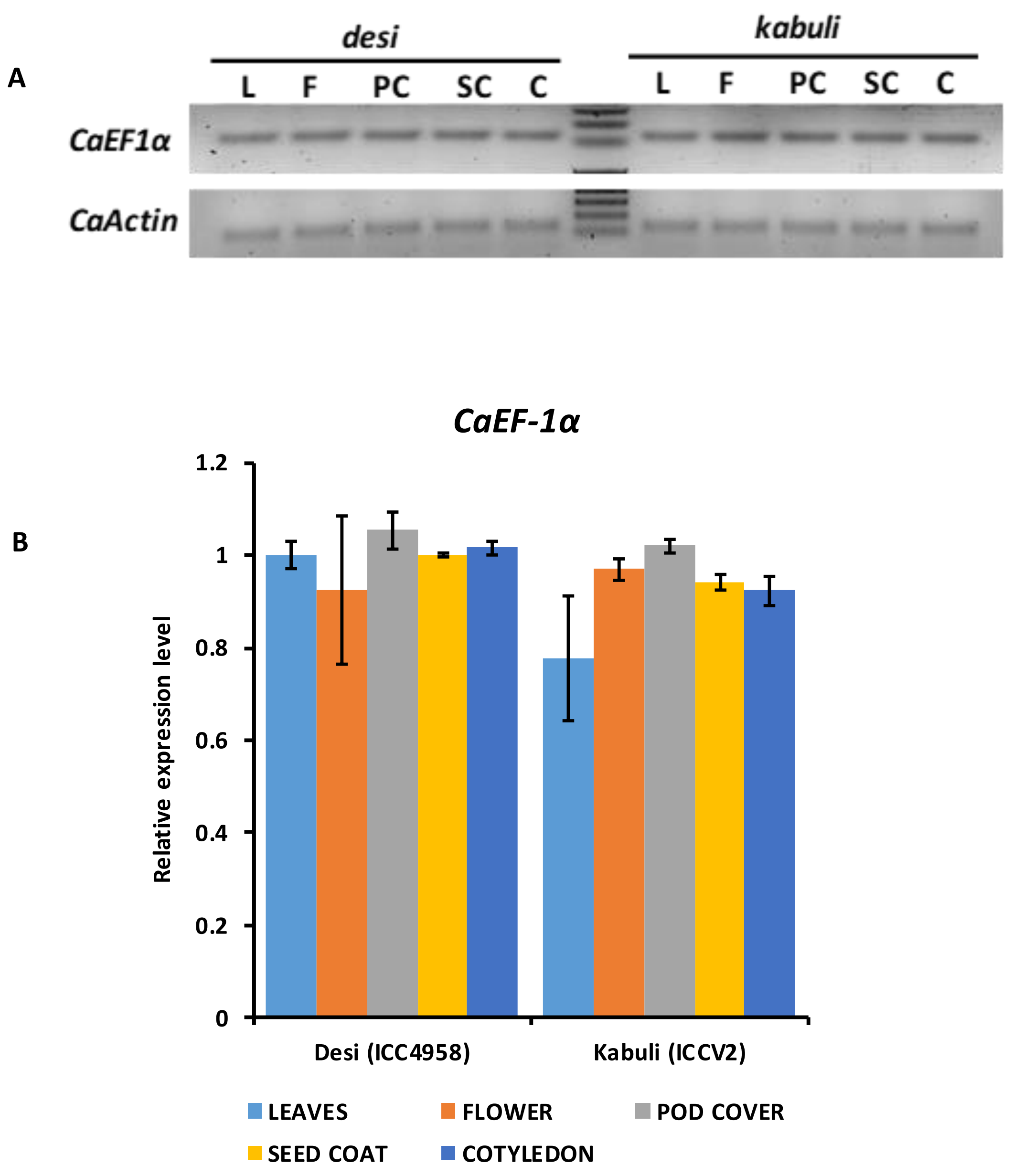


**Supplemental Fig. S1 RT-PCR and relative expression analysis of *CaActin* (AJ012685. 1) with *CaEF-1α* (NM_001365163.1) as endogenous control in different tissues of *desi* and *kabuli*. A)** RT-PCR of *CaEF-1α* and *CaActin* in all the chickpea tissues at 28 PCR cycles. **B**) Relative expression analysis of *CaEF-1α* with *CaActin as* endogenous control. All the assays were performed using three biological replicates, and the bars represent means ± SE.

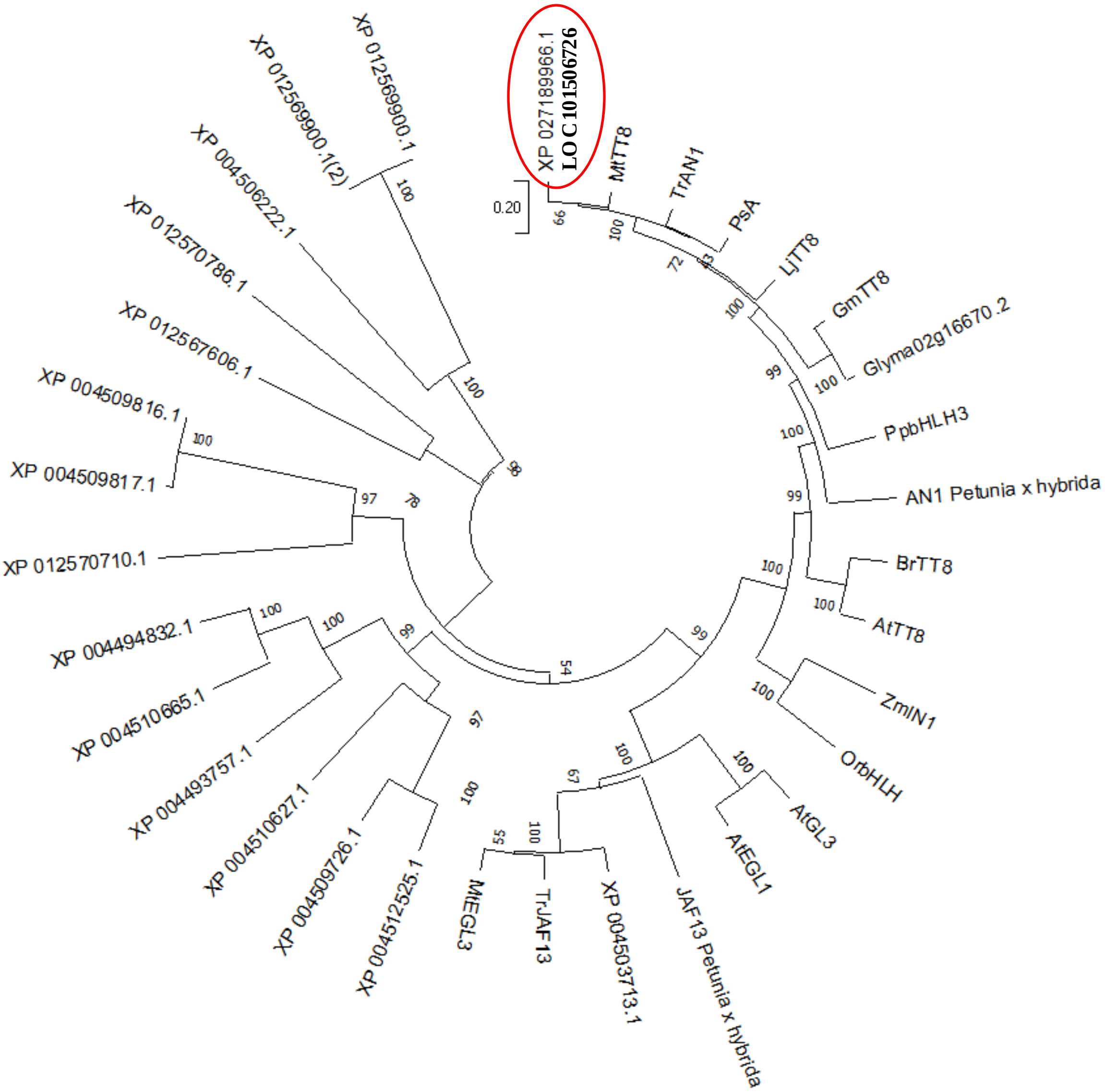


**Supplemental Fig. S2 Phylogenetic tree of basic helix loop helix (CabHLH) and its homologues in chickpea.** The resulting aligned sequences used to the MEGA software (version 10.2.6; http://www.megasoftware.net) (Tamura et al., 2007) by the neighbor-joining method with bootstrap analysis of 1000 replicates. The following protein accession Ids were used; CabHLH (XP_02718996.1, LOC101506726) in red circle; and other bHLH family members (Denoted with XPs); *Petunia x hybrida* PhAN1 (AAG25927.1), GL3 (Q9FN69), EGL3 (Q9CAD0); *Pisum sativum A* (E3SXU4); *Petunia x hybrid* PhJAF13 (AAC39455); *Trifolium repens* TrJAF13 (AIT76563), TrAN1 (AIT76559); *Medicago truncatula* MtEGL3 (Medtr8g098275, KEH21065); MtTT8 (KM892777); *Brassica rapa* BrTT8 (AEA03281.1); *Arabidopsis* TT8 (Q9FT81); *Lotus japonicus* LjTT8 (BAH28881); *Prunus persica* (peach) bHLH3 (AIE57508). *Zea mays* IN1 (AAB03841); *Oryza rufipogon* OsTT12 (ADY17841.1); *Glycine max* TT8 Glyma10g03140.2 (XP_006588626); Glyma02g16670.2 (XP_006575074).

**A**

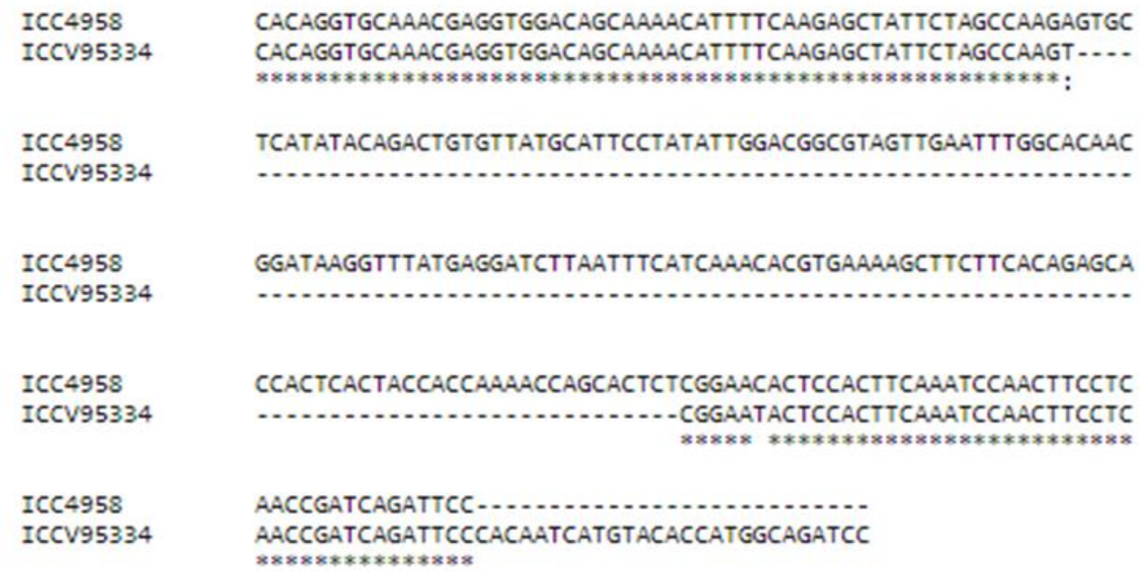


**B**

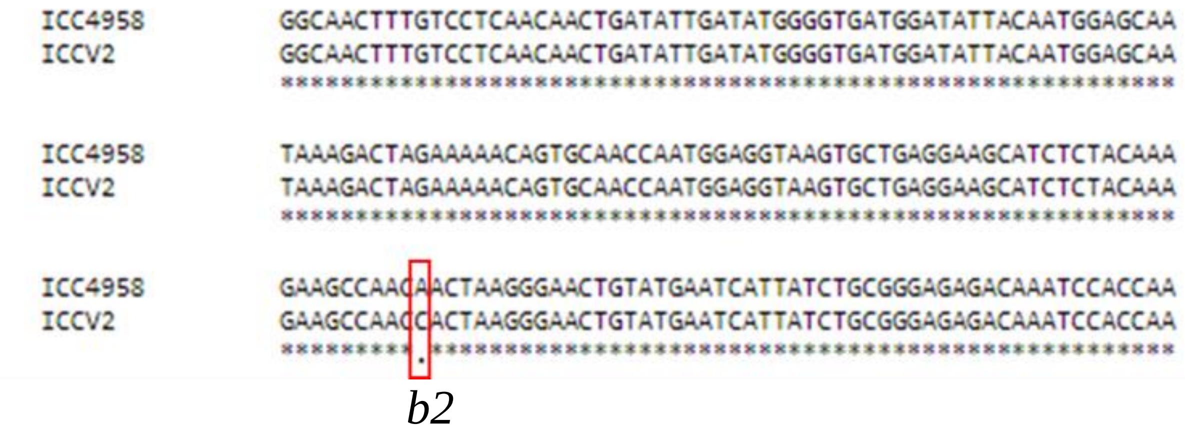


**Supplemental Fig. S3 Alignment result of *bHLH* gene using Clustal W tool. A)** Alignment of ICC4958 (*desi)* and ICCV95334 (*kabuli),* indicates *b4 bHLH* natural allele. **B)** Alignment of ICC4958 (*desi*) and ICCV2 (*kabuli)* indicates *b2* natural allele of *bHLH*.

A

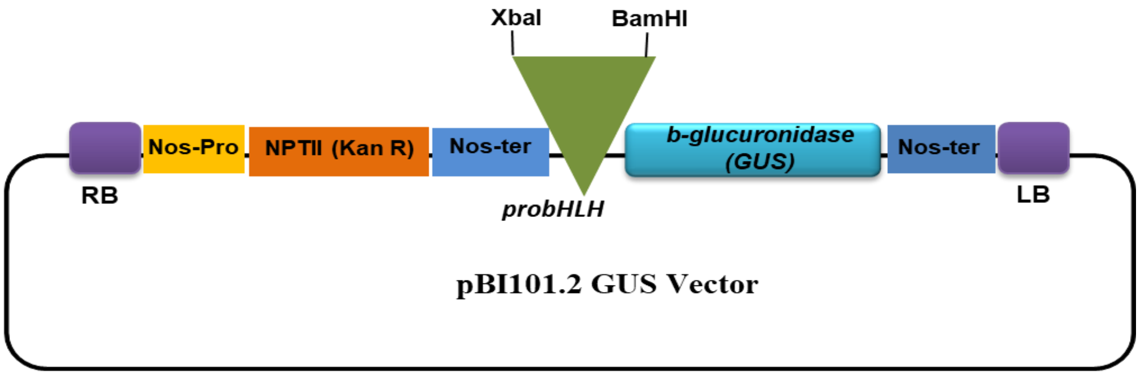


B

CAAACACAACGATTGACACAGACATGTTAAGATTCAGTTCATTTTTTTTTCTTTTAATTTTGTAAATAAATA
TTTATTTTAAAAAAATAGCACTTTTGTAATACAAAATCGGATTTACGTTATTAACTAATTAAAGTACTTCATT
TACTATTAAAAAAATATATTTTAACCACTTATTAAATATACATATATTTCATTAATTATGAAATTATATTATTA
TAAAGTAGTTAATCTTATTCATTTTAATAGTTAATGAGATATATGTTAAATAAATAGTTAATTACATGCATTT
TAATAATTAATATTATATATTTTAATAGTTAATAATATATATGTTAAATAAATAACTAATTACATACATTTTA
GTGGTTAAAGAGTAACTAATGAGATAAACTTATATATGTGTCTTGAGACGGATTCGGAAACAAATTATACT
GGAAATCGACAAAAATTATTATTAAATATTGAATAAACGTAATTTAATAAATATATATTAATTAAAAAATTT
AATTTATAAGTATTGATCATACATCATCATTTAGATTTTAATATATGCATTTTTAAAAAATAATAATATCACG
ACACATTTAGTTAACAAAATTTAATTAAACACTTTTGTACGATGTTTTTAAGCCGAAAGTACATATAATTAA
ATTAAATTCTATTGTTAATATATATGTATATATGTATACTGTGTTATTGGAACATAGACACCCACCGTATATA
TATAATGTATTGTTAATTTTTTTGACAAAAAAAATTAATAAAAAAAGTTATTTTCATATAAAAAAATATAAA
AATATATTTGCAATCCATTATACATATTTGTGTCACTAAAATGCATCTCATTAACCACTTTTTAAATATATAAC
TTATTAACTACTAAAATTAGAATATTTTTTTAAACACTAAATGACATAACTCTAAAGTGGTTAATATTTTATA
TTTTAATAATTAATGAGATATATGTTAAATAAGTGATTAATAATTTATATTTTAACAGTTAATCAAATATTTA
AAGTGATTAAAGACACAAATATGTATATTGTGTCAAAAAAGTGTCTCAATATATAAATCCATTAAACAGTA
CATATGGTGGATATATTGTGTAAAAAGTTTGTGTAATTTTTATAAAATAAAATAAATCTCTGGCTGTTTGGT
GTTTCACTGTTGAGCTTGCGATGGAAATTTGCAAATTGCAAGCAATAGACATTATTTGAACGTGGTACAATA
AAAAACCACGCTGGCGCTCTGTAAAATAACACACAACTAACAAACAACAAACACACTTTGAAATGTGTATA
TATATATAGCACCAAAAACAGTCTGGTAGACCAACCAGTCAAAAATCTTTCTATTTCTTCGTTTCTATTTCTTC
ATTTCAATTCCTTCTATATTTTCTATTTCTTTGGTCTTTACATAATCAACAAACTTTCTTCTCTTTTTTAATTCTC
TTTCTCAACCTCACCTACAAGTCTACTTCTTTTCTTTGCTTTGATTTTTCGTTGATAAGAACCTAACAACAACA
ACAACAAAAACAACAACAAAATACTCTTTTTTTAATTGTCTATTTACTTGTCATGAAACCACAGAAGCCAATT
TTTGGTGTTTCCTACTTTTTTCCTATTTTTGGAAAAGTGATTTCAAGATAAGATTGCAGAGAAATAAAGGAA
AAGCCTGCTGAGTGCTGATGAAAAGCCCGTTTAGGAATTAACA

**Supplemental Fig. S4** ***In planta*** **promoter expression study in chickpea**. **A)** *pBI101.2* plant binary vector harbouring the *PromoterCabHLH::GUS*. **B)** Sequence of 1717 bp promoter used in this study.

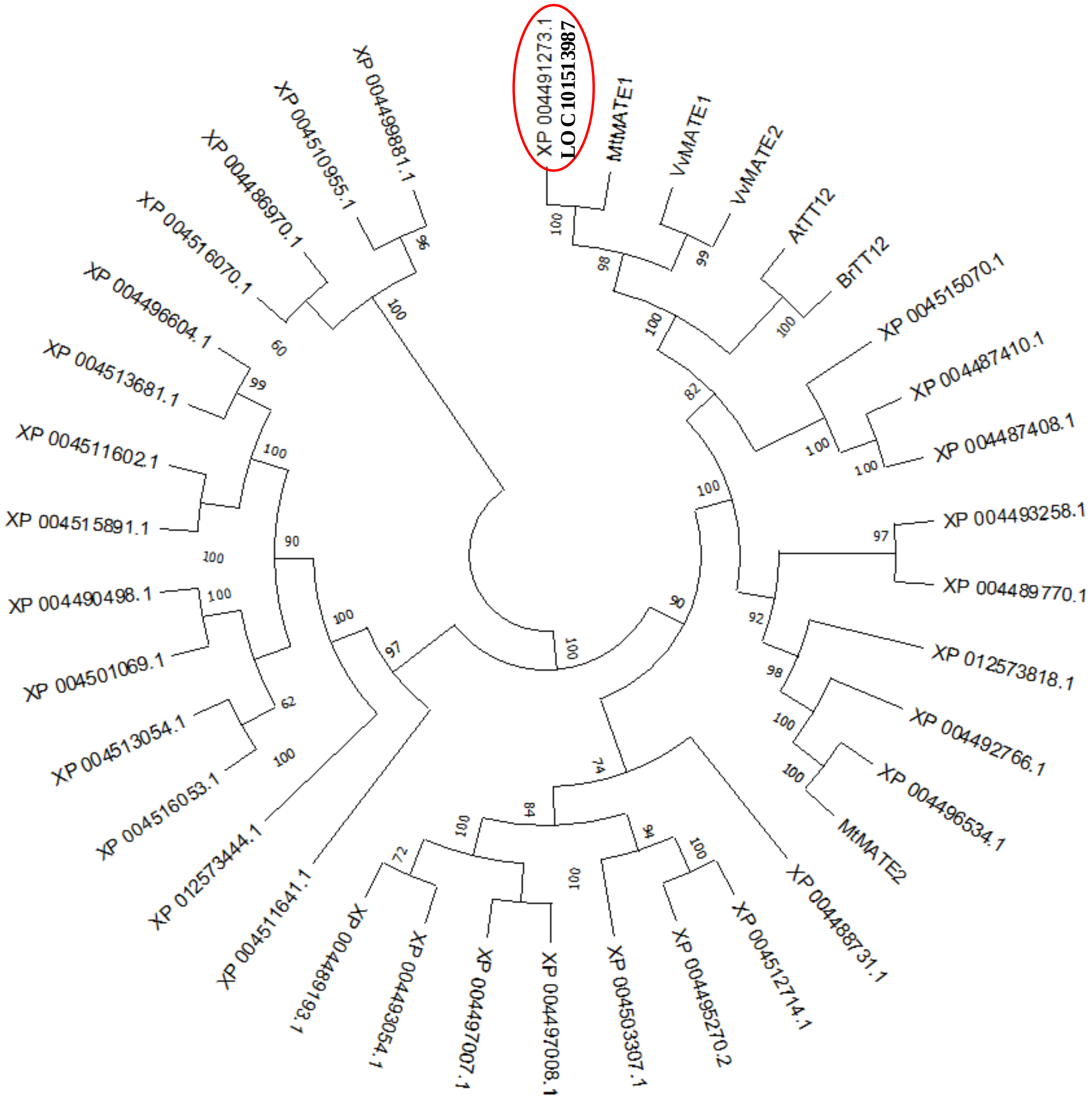


**Supplementary Fig. S5 Phylogenetic tree of Multidrug and toxic compound extrusion (MATE) proteins with other family members.** The resulting aligned sequences used to the MEGA software (version 10.2.6; http://www.megasoftware.net) (Tamura et al., 2007) by the neighbor-joining method with bootstrap analysis of 1000 replicates. The accession numbers were Chickpea MATE1 (XP_004491273.1; LOC101513987) in red circle; and other MATE family members (Denoted with XPs); *Medicago truncatula* MATE1 (ACX37118) and MATE2 (HM856605); *Arabidopsis thaliana* AtTT12 (NP_191462); *Brassica rapa* BrTT12 (ACJ36213); *Vitis vinifera* MATE1 (XP_002282907); MATE2 (XP_002282932).

A

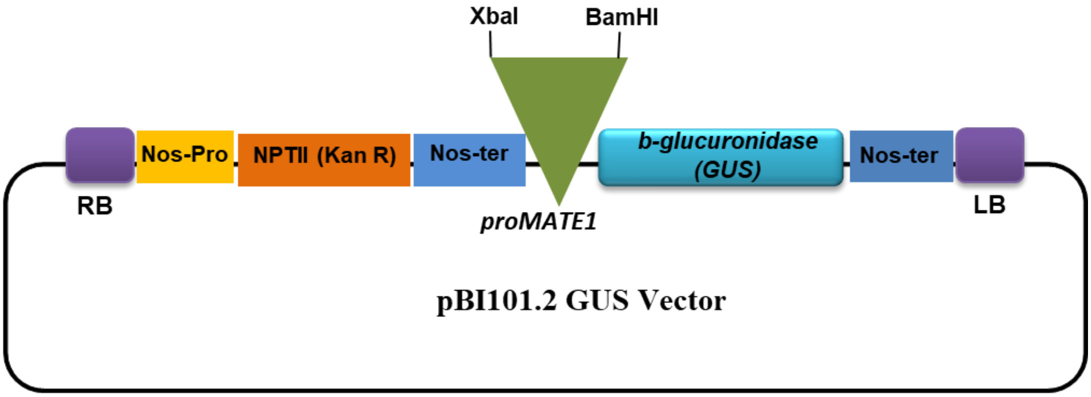


B

GTGTCCGAGTTTCAATAAATAAGATCTAAAGTATATGGTCGATTTTAAATATAAGCATTGTTTTAGAAAATT
AACTATTAATAAAAAAAAATAAAAAAAATAAAAAAATAAATAAAAAATTAGGTAAAAGATATATTTTTAAGGA
TTAAATTTGAATGAGATTAATTTTATAGAAATTAAAAATATATTTTATTTTAATAATAAAAATATTTGTTTTT
ATTAATACAATAAAAGAAATATTAATATGTATCTTTAAGTCACTTGTTATTGTATTAAATATAAATTTTGTAT
CGAAAATTATATTTTTTTACTACAAAATTATGTATTCTATTTTTTATTAATTTTGATTTGAAAAAAAAAAAACT
TTTATACTATGTATTCTATTTTTTCACATGTTCTTTTGTTTACGTTATAACAATGTGGTAAAAAATTATAAATT
TCAGGATTGGAGTTGATAATGGTGGATATGAAGATCAAGGAGTTGAAAACCTGAGATGTAATTTGTTTGC
TTATTAATGTCACATCTCTTCCACGAAGATCAAGTGGTTCTCTAGGCTGGCTAAAGATCAACAAATATTGCT
ATCTGGTTGATGTATATAAGAAGTGAAATATGTTTGTACTTTTACATTTGAAATAGGATTCCCGAGTCAATG
GTTCATTACAGCACTCTTAAGAAAAGTTTTGTAGACCGTTTTCTTTTTGGCTGTAAACTTGTAAAAAAGTTTG
TTATATTTATTGTTCAAATTGTTCGATCTATGTATATTTATATTCTCAACTCCAATGGGATGGACCAAAACCA
AATTATAGGAATCTAAGTCGCCGACATGTCACAAATAACTCTTTGAAAACTTTTTTGTACGATAGTTGCTTA
TACTAGTATTTTATTTTTTATTTTTTTCTGATTGAAGCTTGTTCTAGTTTTTTTTTTTTTTTGTTTATAATATCTT
ATACTAGGATTTTGATAGACTTCAGCAAACTTATGTACATTACTGACTTCAAGTTAATTATTATACACTAATT
TTGGGCCAGATGCTCTTTCAGCTCCAGGATTATATTATAGACTTTTTACTGAATCCCATGATGATTCAGAAG
TGAAAATGTCGATCCGACAGTGAAAAATGAGATTAATGAATTTACCAAAAGTTCTAATTTTAACATATTTTA
AATAAACATTGAGGATGAATGAGCAACCACTTAATCTTAATCTTCGTGCTATTTAATGATCTCTATTGAACC
AATCTCTTAGGTCAAGAACAAAATAAATATTTTAAATATTAATATTTAATTTTCTTCGAAATCTGAAGGGTA
TTAAATATTATTTTAAATAAACATAAGTAGCATATTTCTATGCGGTAATTCTTGAACACAAGCTCACTCATTT
ACCATAACACAAATCATTTAATTTCGGCCATAATATGCTATGGTTCAGGTGAGAGATTAAGACATTCTAAAA
TGTGGGCCTAAGAAGTAACATGTTAGAGGCATTAATGTACCCCTAACACACACACAAGAGCAATGACTCAA
CTAAAAGTCTTATTCACAACTAAAGGTCAGTGCTAGTCATTTTCAACAAGTTAGATTCTAGGAGAGACCAA
AAAGGTTTGTGTGAGAGGGAATGAATGAAGTGGTGCTATTCTCCCAACTTTTGAATTCAAATGAAACTCAT
TGTTATAAAGGAAGAAAAAGGGCTAGAAGGGGTGGTAGTTTCATCTTCTTGCCTATGTTTAGAGGAACGT
GAATAAGAGGGGTAGGTAGTAATTAATAACTATCAACAACAAAAATAAAATGGAAAACAACAACAAATCA
AAAAAATATCCTATTGCCAACAACAAAATGTGGAAGGGTAGCTACAAAATTAAAAGATAGAAAAAATCTC
ATTAATAATGTTCTACTTGTAATGTTAACGCTTCATTCCCATGCATTGAACTAGTTGTGACTTGTGAGGCACA
TAACC

**Supplementary Fig. S6 *In planta* promoter expression study in chickpea**. **A)** *pBI101.2* plant binary vector harbouring the *PromoterCaMATE1::GUS*. **B)** Sequence of 1950 bp promoter used in this study.

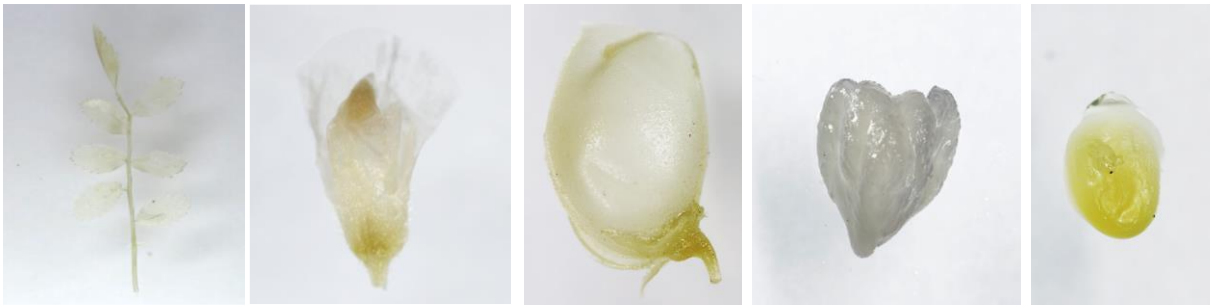


**Supplementary Fig. S7** ***In planta*** **visualization of no promoter GUS (*β-glucuronidase*) activity in various tissues of chickpea.** Leaves, flower, pod cover, seed coat and cotyledon of T2 transgenic plants transformed with a GUS construct without any promoter did not show GUS-stain.

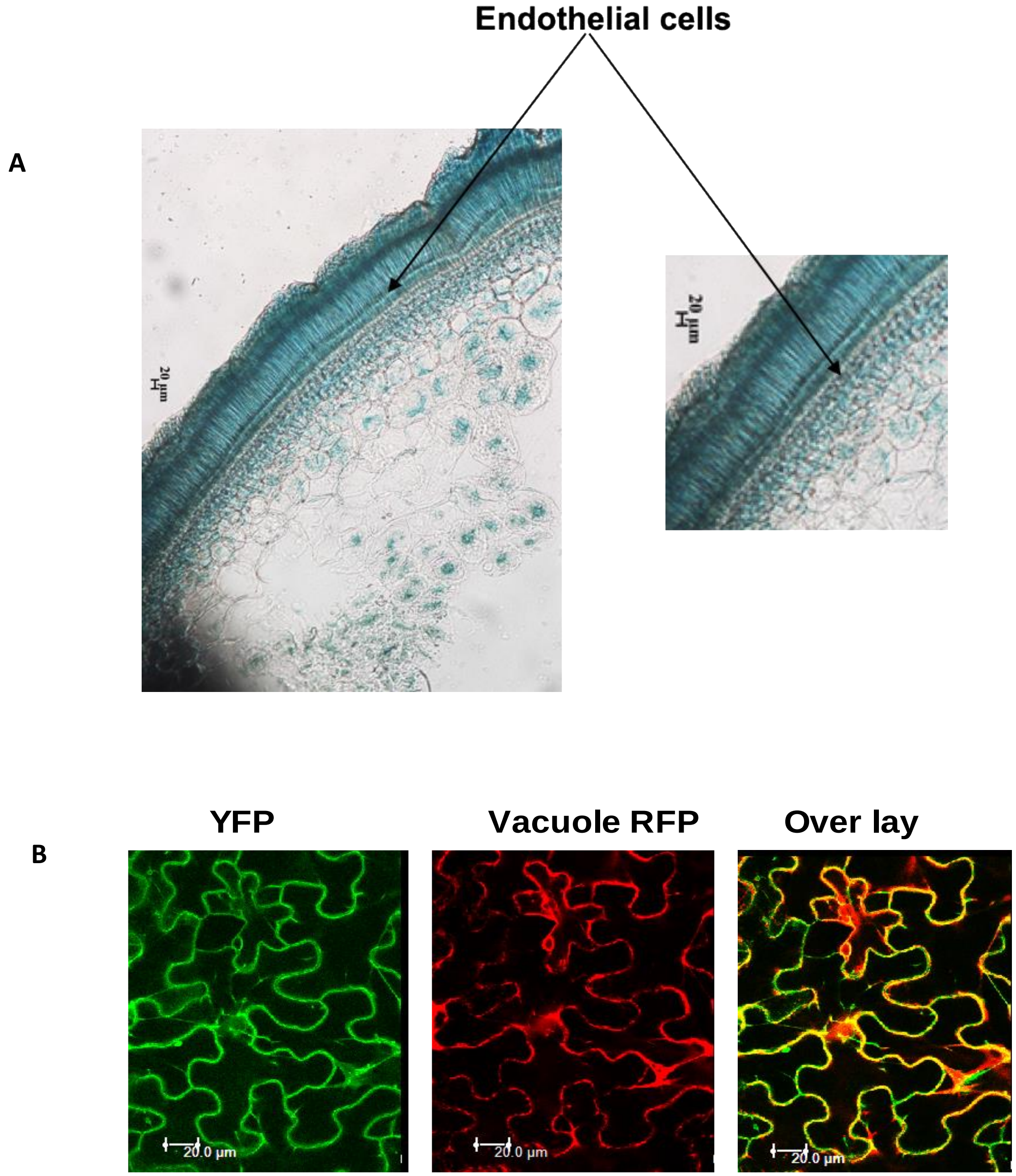


**Supplementary Fig. S8 Cross-section of *PromoterCabHLH::GUS* construct transgenic seed lines and subcellular localization of CaMATE1 protein. A)** Cross- section of seed coat showing *in planta* visualization of GUS (β-glucuronidase) activity of Promoter*CabHLH*-GUS. Scale bar, 20 µM. **B)** Subcellular localization of CaMATE1. *CaMATE1-YFP* along with vacuolar membrane (VM) marker (CD3-975 mCherry) was co-agroinfiltrated in *N. benthamiana* leaves and was visualized under confocal microscope after 60-65 hrs. Scale bars, 20 µm.

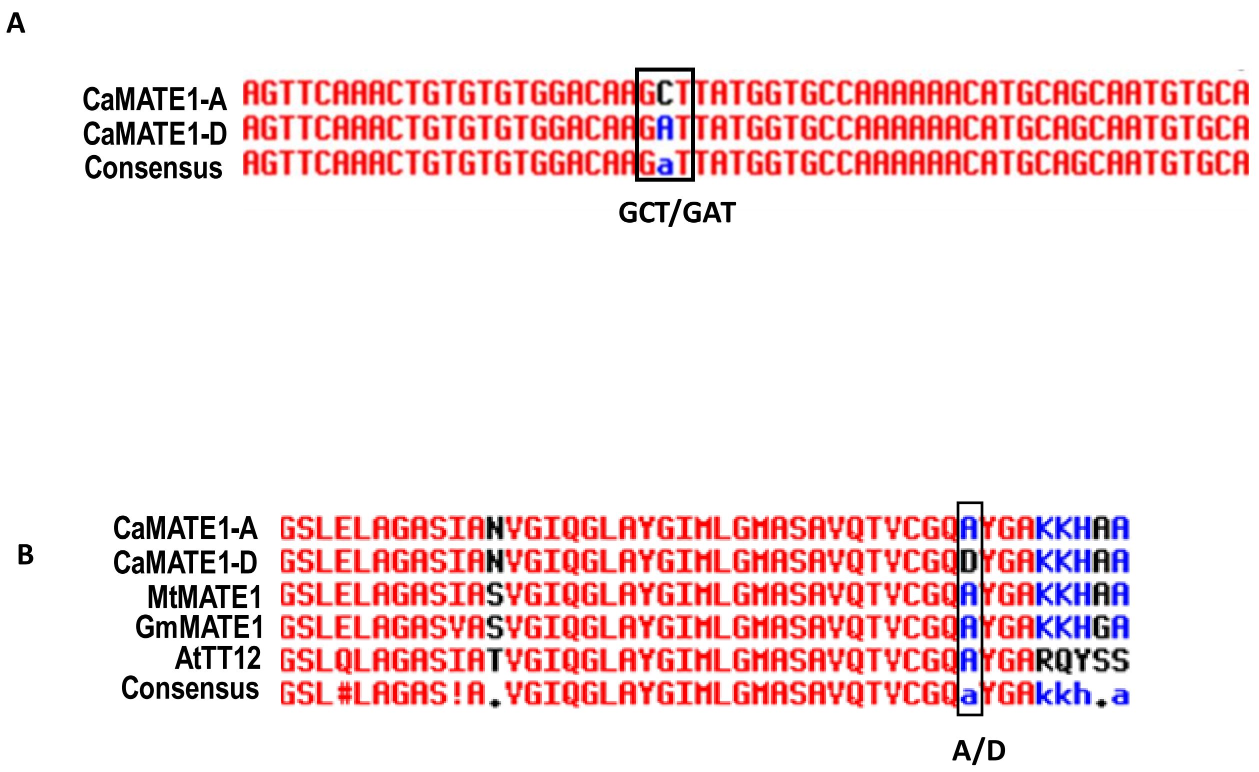


**Supplementary Fig. S9 Sequence alignment of MATE1 from different plant species**. **A**) Nucleotide sequence aligned of CaMATE1-A and CaMATE1-D. CaMATE1-A and it's variant CaMATE1-D were shown separately. **B**) Alignment of protein analysis was done with Multialign tool. Black box indicates the conserved amino acid Alanine and it's variant Aspartic acid (D) in chickpea. Protein sequences and their accession numbers are; *Medicago trancatula* ; MtMATE1 (XP_024639756.1), *Glycine max*; GmMATE1 (XP_003545155.1), *Cicer arietinum* ; CaMATE1 (XP_004491273.1), *Arabidopsis thaliana* ; AtTT12 (NP_191462.1). The position of the CaMATE1 allele has been marked (A/D).

A

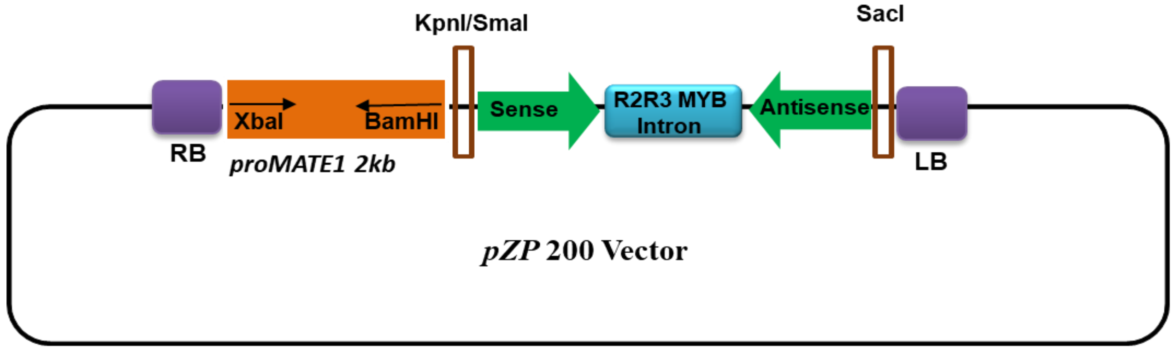


B

GTTATTGTTGGATGTTATGCAAATGCTTAGGGAGATGAGAATTGAGGTTATTGGTGTTCAATCTTCG
CTTAACAATAATAATGGTGTCTTTGTTGCGGAATTAAGAGCTAAAGTTAAGGATAATGGAAATGGG
AAAAAAGTTAGCATTGTTGAAGTTAAGAGAGCACTTAATCAAATTATTCCTCATGCCAATTACTAGCT
TAGTACGGTGTTGCACTTGTGCTGATAATTAGATTTTAAAAGATAAATATTAAATATTTTGATATTTA
TCTCTTTGAATCTAATTGTCTCTGTAGTTGATGTGCGTATGGATCCTCCGGTAATCATCATTCTAGTTG
TCTCTTTTTTTTTATTACCGCAAACAAAATGTTATATAATCTATATATACATCATATATTAAAAAGTAC
AACCTGAGAAATAAATTTAACGATGTTTCTAAAGTGAAGTTTTACTTCGAAAAAATTTATTCGTTCTT
TTTTCCAATGTATTTGACAATTCTCTGATGCAAAATATGTGTTTGGTTAGCACACATCATTTAGTCTAT
ATTCCATAAAAAAACTTCAAAATAAATTTTATTAACTGTGGTCTTCCATCTTATCTCTTTCACTATTCTT
GTCTTTAGGTGGTCGGCAACTACAGAGACAATTAGATTCAAAGAGATAAATATCAAAATATTTAATA
TTTATCTTTTAAAATCTAATTATCAGCACAAGTGCAACACCGTACTAAGCTAGTAATTGGCATGAGGA
ATAATTTGATTAAGTGCTCTCTTAACTTCAACAATGCTAACTTTTTTCCCATTTCCATTATCCTTAACTTT
AGCTCTTAATTCCGCAACAAAGACACCATTATTATTGTTAAGCGAAGATTGAACACCAATAACCTCAA
TTCTCATCTCCCTAAGCATTTGCATAACATCCAACAATAAC

C

GGAATGCAGAGGTAATCATCACTTTTGGCCTAGACTAGAATAACTTATCTGTTACGTGTTCCACATGA
ATTGTTTGCCACTTAGGAAGGTGTTAAACGATTTATAATGTTGTTGTAGGTTGAAAAAGCAATTGTTC
GTGTCAGGAGATCTGCTGAAGATGAGACCTTGGATCAACTGGTTTCCAACGTATAGGACATTTATTA
CTGTAACTTTTCTTTTCTAGGAATGGAAGCAGAACACTTTTAGCAGCAGATTAGTCTTGTTTTTGGAC
TATAAGGGATTGTTGTTGGGATGTGCGTATGGATCCTCCGGTAATCATCATTCTAGTTGTCTCTTTTTT
TTTTATTACCGCAAACAAAATGTTATATAATCTATATATACATCATATATTAAAAAGTACAACCTGAG
AAATAAATTTAACGATGTTTCTAAAGTGAAGTTTTACTTCGAAAAAATTTATTCGTTCTTTTTTCCAAT
GTATTTGACAATTCTCTGATGCAAAATATGTGTTTGGTTAGCACACATCATTTAGTCTATATTCCATAA
AAAAACTTCAAAATAAATTTTATTAACTGTGGTCTTCCATCTTATCTCTTTCACTATTCTTGTCTTTAGG
TGGTCGGCCCAACAACAATCCCTTATAGTCCAAAAACAAGACTAATCTGCTGCTAAAAGTGTTCTGCT
TCCATTCCTAGAAAAGAAAAGTTACAGTAATAAATGTCCTATACGTTGGAAACCAGTTGATCCAAGG
TCTCATCTTCAGCAGATCTCCTGACACGAACAATTGCTTTTTCAACCTACAACAACATTATAAATCGTT
TAACACCTTCCTAAGTGGCAAACAATTCATGTGGAACACGTAACAGATAAGTTATTCTAGTCTAGGCC
AAAAGTGATGATTACCTCTGCATTCC

**Supplementary Fig. S10 Schematic representation and Sequence of RNAi Cassette including sense (yellow), R2R3 MYB intron (green) and antisense (blue). A**) *pZP200* plant binary vector used for the development of RNAi construct. **B**) *CabHLH* RNAi construct **C**) *CaMATE1* RNAi construct.

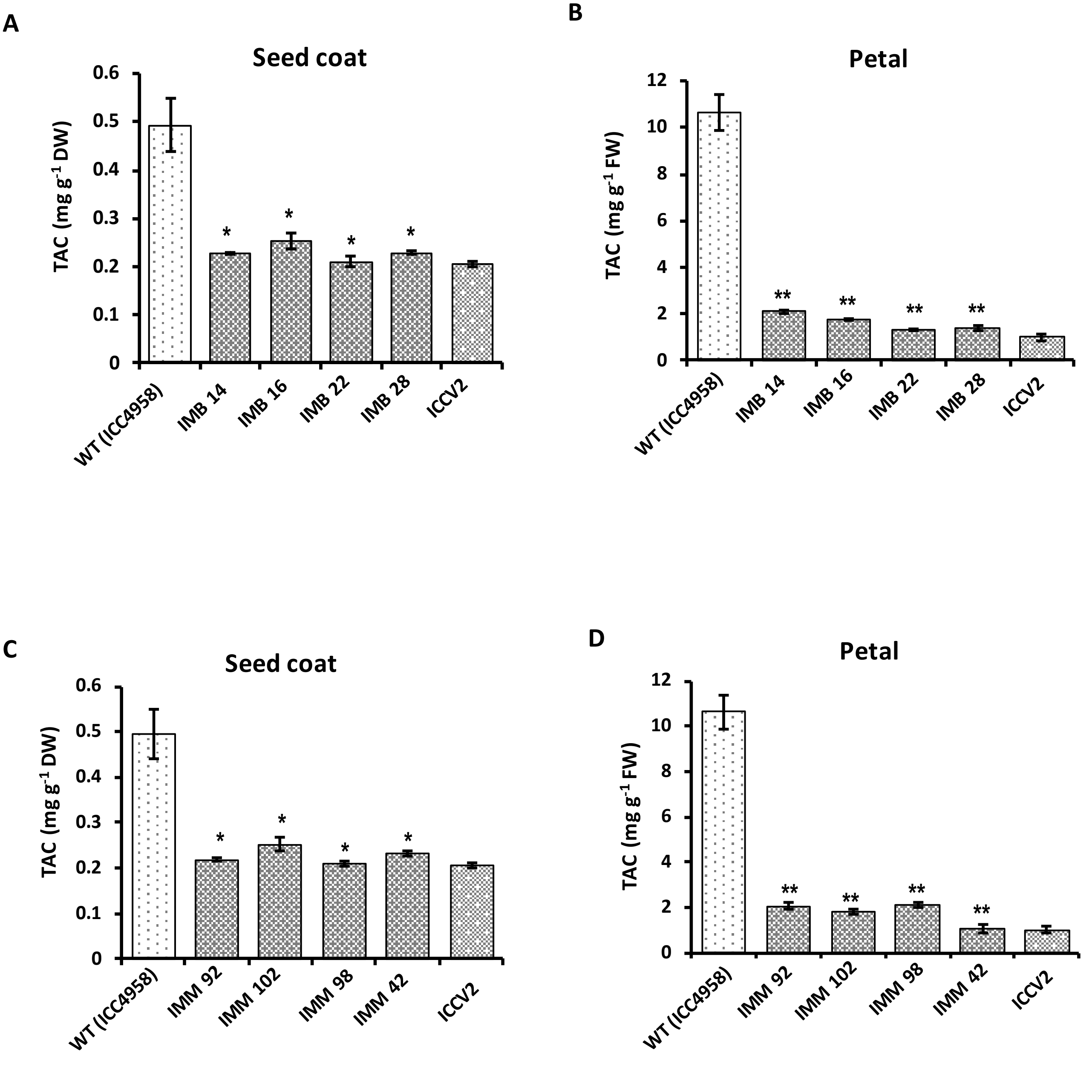


**Supplementary Fig. S11 Estimation of total anthocyanin in flower and seed coat in four independent transgenic lines. A, B**) A total anthocyanin quantification (TAC) in *PromoterMATE1::CabHLH* RNAi transgenic plants of seed coat (left) and petal (right). **C, D**) A total anthocyanin quantification (TAC) in *PromoterMATE1::CaMATE1* RNAi transgenic plants of seed coat (left) and petal (right). DW and FW denotes dry weight and fresh weight respectively. The error bars represent ±SD. Asterisks indicate significant differences from the WT (ICC4958) as determined by two-tailed Student's *t*-test (* for $p < 0.05$; ** for $p < 0.01$).

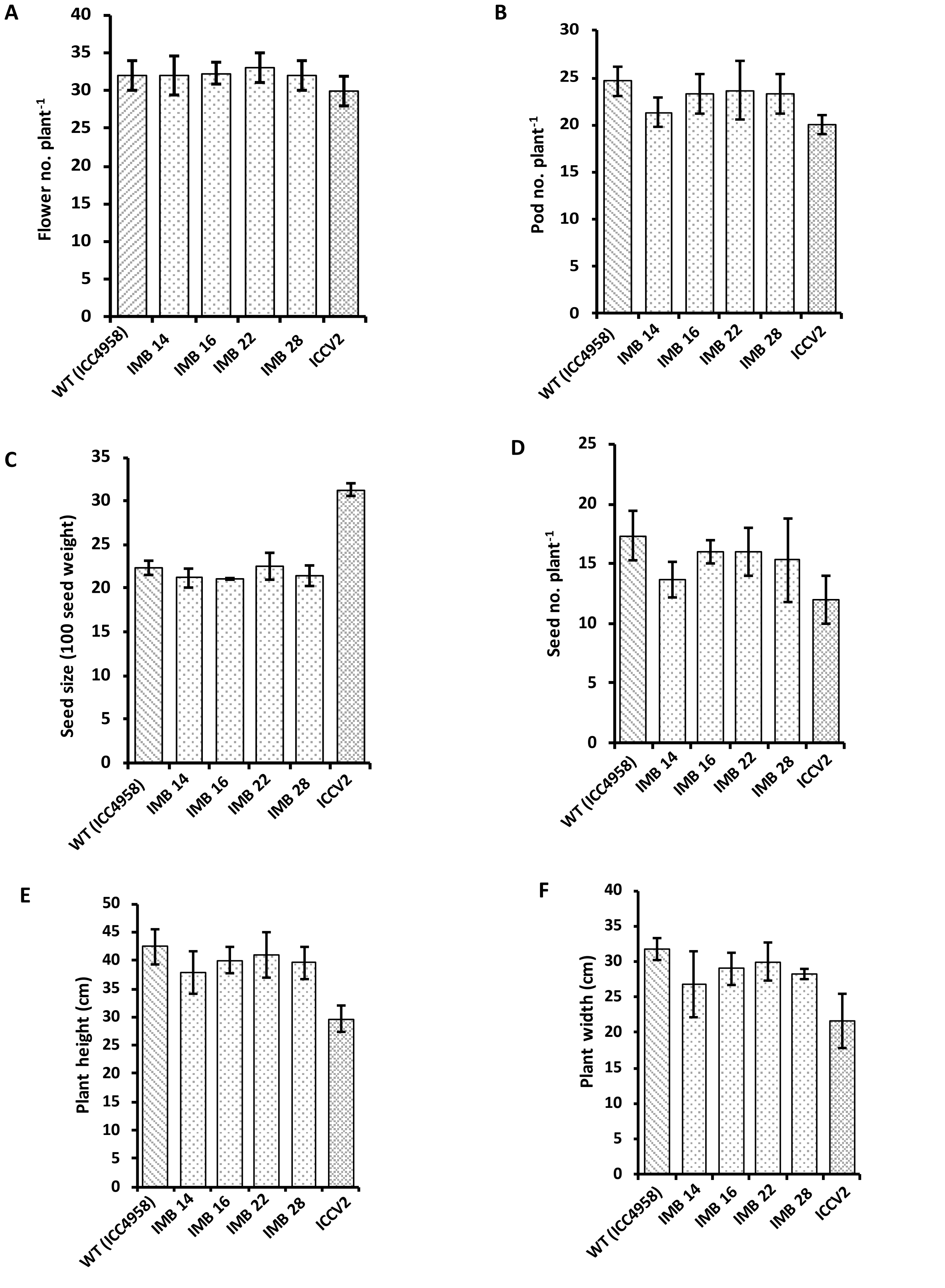


**Supplementary Fig. S12 Comparative phenotypic data for various agronomic trait in transgenic lines expressing the construct *PromoterMATE1::CabHLH* RNAi. A)** evaluation of number of flowers per plant **B**) pod number per plant **C)** seed number per plant **D**) seed size (100 seed weight) **E-F**) plant height and plant width (in cm) . The error bars represent ±SD.

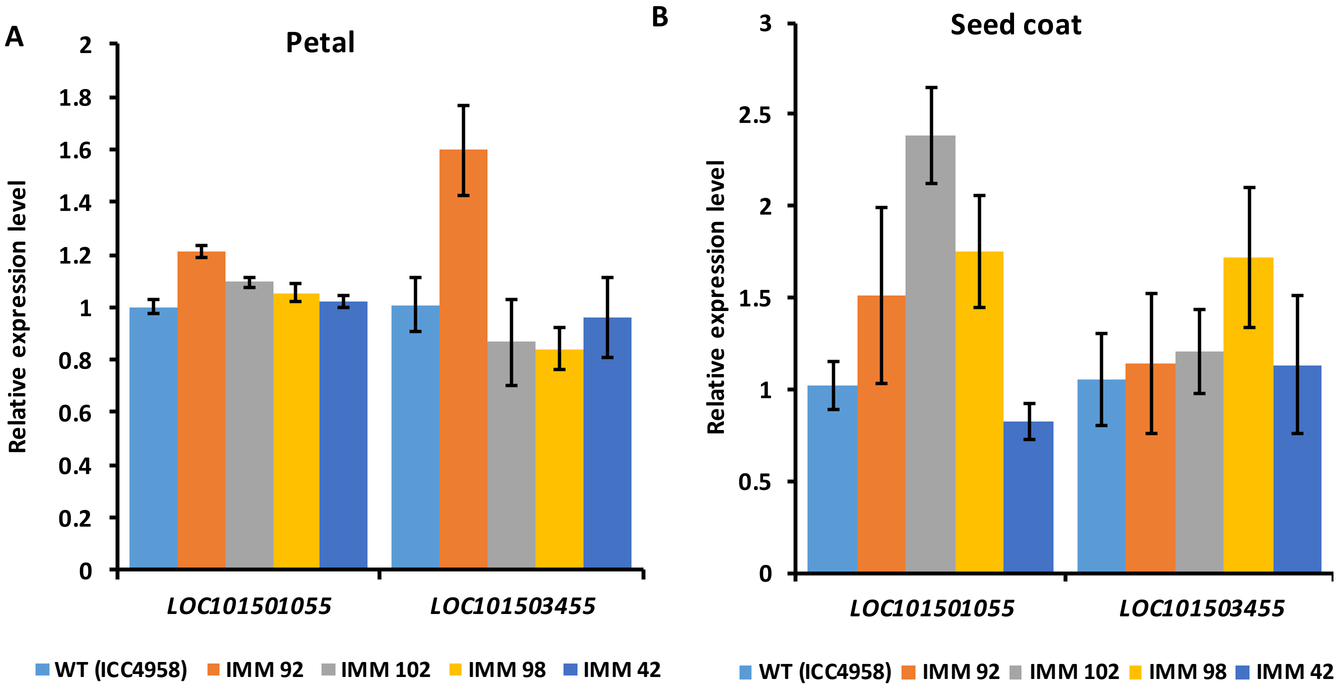


**Supplemental Fig. S13 Relative expression analysis of two *CaMATE* genes (LOC101501055, LOC101503455) in *PromoterMATE1::CaMATE1* RNAi transgenic lines. A-B**) Expression analysis of two MATE genes, LOC101501055 and LOC101503455 genes in four independent RNAi lines and WT (ICC4958) cultivar in petal and seed coats. *CaEF-1α* and *CaActin* were used as the internal control. Bars in represent means ± SE. Statistically significant differences from chickpea accession ICC4958 (WT) determined using two-tailed Student's *t*-test.

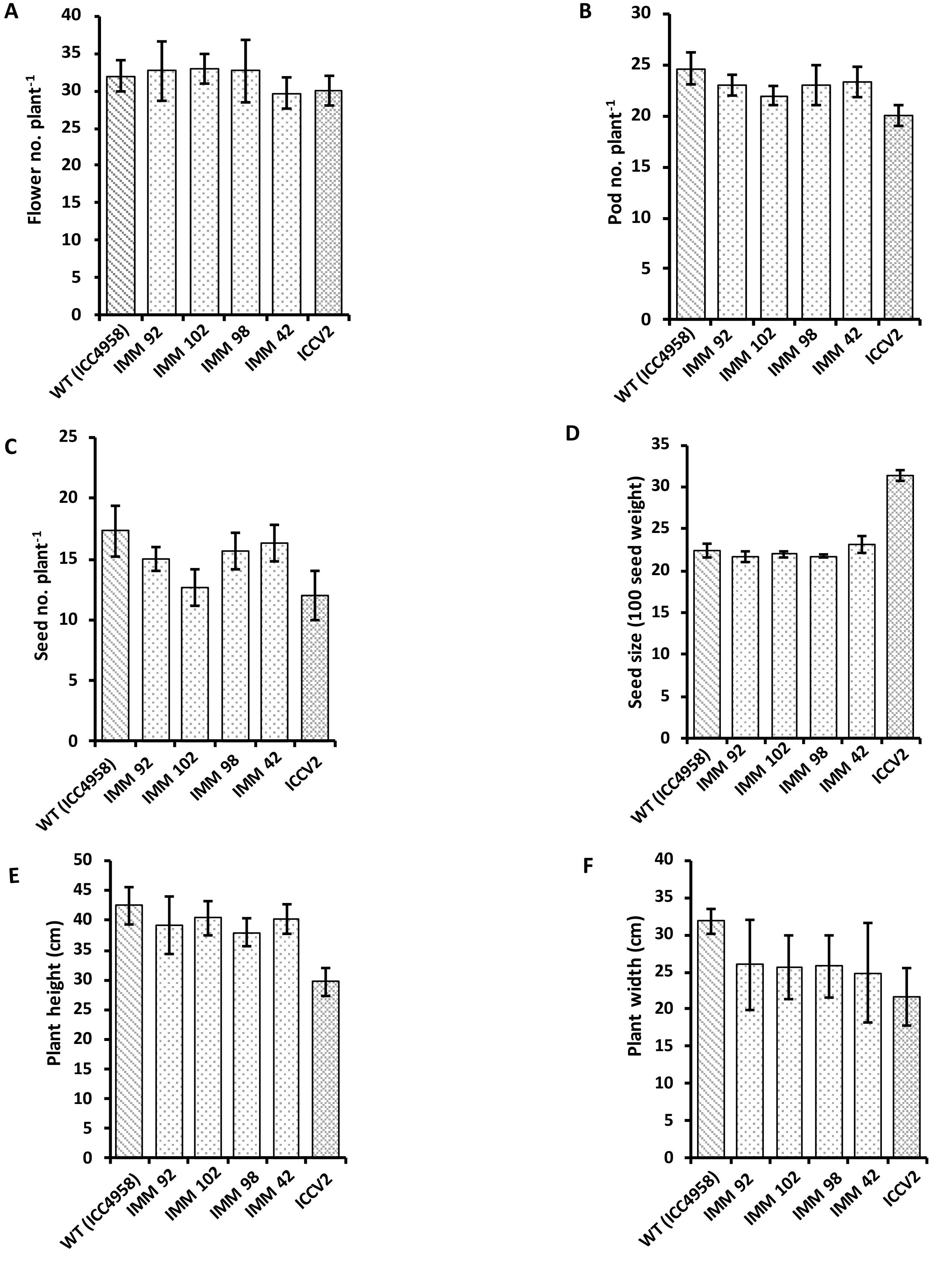


**Supplemental Fig. S14 Comparative phenotypic data for various agronomic trait in transgenic lines expressing the construct *PromoterMATE1::CaMATE1* RNAi. A**) Evaluation of number of flowers per plant **B**) pod number per plant **C**) seed number per plant **D**) Seed size (100 seed weight) **E-F**) plant height and plant width (in cm) . The error bars represent ±SD.

**Supplementary Table 1** Primers used in this study

| Primer name | Sequence (5'to 3') |
|---|---|
| bHLHp_F (Xba1) | CTAGTCTAGACAAACACAACGATTGACACAGA |
| bHLHp_R (bamH1) | CGGGATCCTGACATTGTTAATTCCTAA |
| bHLH SF (NcoI/Sma1) | CATGCCATGGCCCGGGGTTATTGTTGGATGTTATGC |
| bHLH SR (SacII) | TCCCCGCGGCAACTACAGAGACAATTAG |
| bHLH_ AsF (Pst1) | AACTGCAGCACATTTGGCTTGAGATGA |
| bHLH _AsR (Sac1) | CGAGCTCGTTATTGTTGGATGTTATGCA |
| MATE1p_F (Xba1) | GCTCTAGAGTGTCCGAGTTTCAATAAATAAGATC |
| MATE1p_R (bamH1) | GGATCCTGGGTGGTGTTCCATAGTGCTCAT |
| Ca_MATE1_F_SQ | ATTGGGTTGAGATGGTGGCT |
| Ca_MATE1_R_SQ | GAGAAACACTGCTGCACCAA |
| MATE1_SF(NcoI/Kpn1) | CATGCCATGGGGTACCAAGGAATGCAGAGGTAATCACTT |
| MATE1_SR (SacII) | TAATCCGCGGCCCAACAACAATCCCTTATAGTCC |
| MATE1_ASF (Pst1) | ACACTGCAGAACCCCCAACAACAATCCCTTA |
| MATE1_ASR (Sac1) | CCGAGCTCCAGAGGTAATCATCACTT |
| Intron FP | ATGTGCGTATGGATCCTCCGGTAATCA |
| Intron RP | CCGACCACCTAAAGACAAGAATAGT |
| Seq_F1_bHLH | GGCAACTTTGTCCTCAACAACTG |
| Seq_R1_bHLH | GATTCTGTTAAGTCCTCAGGCGA |
| Seq_F2_bHLH | CACAGGTGCAAACGAGGTGGAC |
| Seq_R2_bHLH | CGTCGTCTTCGTCCATGTCATCT |
| Seq_F3_bHLH | AGATCATTGGTCCCTTTTGTTAC |
| Seq_R3_bHLH | CTATTTGACGGTTACGCGTTTCG |
| qRT_bHLH_F | TGCTTCCTTGTCGCCTGAGGACT |
| qRT_bHLH_R | GCAACCCGACACCAGGAGGAAAA |
| qRT_MATE1_F | GGCATATATGGCAGTTGGAGTA |
| qRT_MATE1_R | GCTGCACCAAGAAGGCCATAG |
| qRT_DFR_F | GAGAATCCAAAAGCTGAAGGGAG |
| qRT_DFR_R | TTTGAATCCCAACTCAGTGATCT |

| Primer name | Sequence (5’to 3’) |
|---|---|
| NPTII_F | ATGATTGAACAAGATGGATTGC |
| NPTII_R | ATGGGTCACGACGAGATCATCG |
| MATE1pE_F CDS | CACCATGAGCACTATGGAACACCA |
| MATE1pE_R CDS | TACGTTGGAAACCAGTTGATCC |
| bHLHpE_F CDS | CACCATGTCAATGGCTGCTCCACCAGTTG |
| bHLHpE_R CDS | GTAATTGGCATGAGGAATAATTTG |
| qRT_BAN_F | CTGAAAAGGCTGCATGGAA |
| qRT_BAN_R | ATTGCCAAGCCAACACTAGC |
| qRT_LDOX_F | AGAAGGGCCTGAAGTTCCAA |
| qRT_LDOX_R | TGCATCACACCCCATTCTT |
| SQbHLH_F | GATCCACCGTCCACAAATCTCAA |
| SQbHLH_R | GTTGGATCCATCGTTTGGTGAAC |
| Ca_PAR qRT_F | CATTAGGCTTCACAAACTATTAGG |
| Ca_PAR qRT_R | CATGTTGGTGTGTTGTTTATGGA |
| CaTTG1 qRT_F | GGCCTTCTCTTCAAACCCCAA |
| CaTTG1 qRT_R | TGGAGAGGGTATCAGGGTTGA |
| Ca_PAR CDS _F | CACCATGGGAAGAAGCCCTTGTTGT |
| Ca_PAR CDS_R | TCATTCTCCTAGTACTTCCTC |
| CaWD40 CDS_F | CACCATGGAGAATTCCACTCAAGAA |
| CaWD40 CDS_R | TCAAACCCTCAAAAGCTGCAT |
| Ca_LAP1_F | CACCATGGGTGGTGTTGCATG |
| Ca_LAP1_R | AGAGGAAAAATCATTATAAAGATC |
| MATE1CDS_F (SmaI) | TTCCCCCGGGATGAGCACTATGGAACACCACCCA |
| MATE1CDS_R(Sal1) | ACGCGTCGACCTATACGTTGGAAACCAGTTGATC |
| MATE1_SDM_F | GTGTGTGGACAAGATTATGGTGCCAAA |
| MATE1_SDM_R | TTTGGCACCATAATCTTGTCCACACAC |

| Primer name | Sequence (5’to 3’) |
|---|---|
| CaActin_F | CCTGAAGAACACCCTGTTCT |
| CaActin_R | GTCTCAAACATGATCTGAGTC |
| CaM455_F | CATAGCAGGCCCATCAATCTTC |
| CaM455_R | TCAGCATCACCTATGCGACC |
| CaM055_F | CACACATCAGTGCCTATGGTG |
| CaM055_R | GTAGAGTTCACCTAAGTGACC |
